# Differential KEAP1/NRF2 mediated signaling widens the therapeutic window of redox-targeting drugs in SCLC therapy

**DOI:** 10.1101/2024.11.06.621846

**Authors:** Jana Samarin, Hana Nuskova, Piotr Fabrowski, Mona Malz, Eberhard Amtmann, Minerva J. Taeubert, Daniel Pastor-Flores, Daniel Kazdal, Roman Kurilov, Nicole de Vries, Hannelore Pink, Franziska Deis, Johanna Hummel-Eisenbeiss, Kamini Kaushal, Tobias P. Dick, Gerhard Hamilton, Martina Muckenthaler, Moritz Mall, Bryce Lim, Taishi Kanamaru, Glynis Klinke, Martin Sos, Julia Frede, Aubry K. Miller, Hamed Alborzinia, Nikolas Gunkel

## Abstract

Small cell lung cancer (SCLC) patients frequently experience a remarkable response to first-line therapy. Follow up maintenance treatments aim to control residual tumor cells, but generally fail due to cross-resistance, inefficient targeting of tumor vulnerabilities, or dose-limiting toxicity, resulting in relapse and disease progression. Here, we show that SCLC cells, similar to their cells of origin, pulmonary neuroendocrine cells (PNECs), exhibit low activity in pathways protecting against reactive oxygen species (ROS). When exposed to a novel thioredoxin reductase 1 (TXNRD1) inhibitor, these cells quickly exhaust their ROS-scavenging capacity, regardless of their molecular subtype or resistance to first-line therapy. Importantly, unlike non-cancerous cells, SCLC cells cannot adapt to drug-induced ROS stress due to the suppression of ROS defense mechanisms by multiple layers of epigenetic and transcriptional regulation. By exploiting this difference in oxidative stress management, we safely increased the therapeutic dose of TXNRD1 inhibitors *in vivo* by pharmacological activation of the NRF2 stress response pathway. This resulted in improved tumor control without added toxicity to healthy tissues. These findings underscore the therapeutic potential of TXNRD1 inhibitors for maintenance therapy in SCLC.

**Graphical summary:** 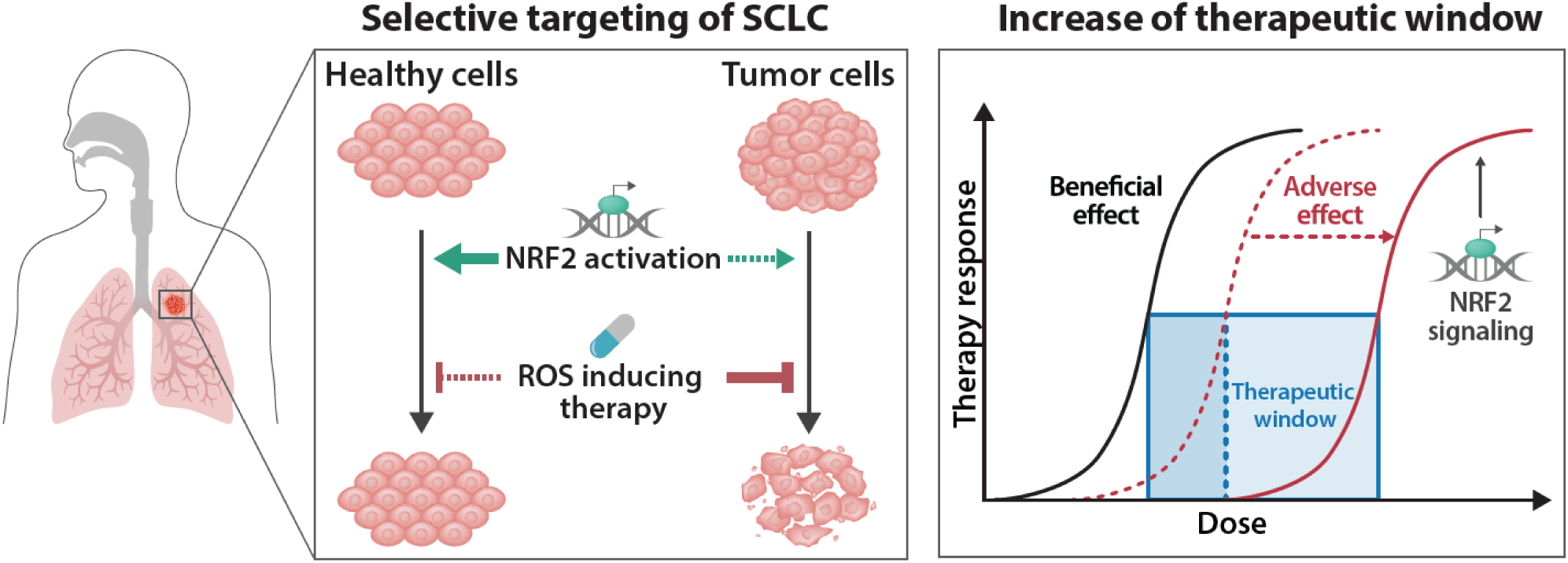

Pharmacological induction of NRF2 leads to differential cyto-protection against TXNRD1 inhibitors in normal tissue but not in SCLC tumor cells. This results in a reduction of adverse effects, allowing to increase the therapeutic dose.

## Introduction

Small cell lung cancer (SCLC) accounts for approximately 15% of all lung cancer cases and is considered one of the most lethal cancer types, with a 5-year survival rate of less than 7% [1]. The main challenges in SCLC include rapid tumor growth, early metastatic spread, and a tendency to relapse with profound therapy resistance, resulting in limited treatment options. Standard-of-care (SoC) treatments target the exceptionally high proliferative rate of SCLC cells and their dependency on the biosynthetic pathways required for cell replication. This is achieved through the use of DNA cross-linking agents like cisplatin or carboplatin, topoisomerase inhibitors such as etoposide or topotecan and γ-radiation, all of which target DNA synthesis, replication and repair [2–4].

Given that 70-80% of SCLC patients respond well to chemotherapy, it seems reasonable to assume that maintenance therapy represents a promising strategy. However, the success of maintenance therapy depends on the vulnerability of the treated tumor, its inability to develop cross-resistance mechanisms during the cytotoxic assault by the drug and conversely, the presence of robust defense mechanisms in healthy tissue to mitigate the drug’s side effects. Due to this complex set of requirement, targeting cell cycle control via ATR, WEE1 and CHK1 inhibitors, as well as DNA repair pathways through PARP1 inhibitors such as olaparib and veliparib has failed to provide substantial benefit for patients [5]. More recently, a combination of SoC and immunotherapy has been explored in clinical trials, resulting in a transient delay of tumor relapse for a few months [6]. Another promising approach is the inhibition of LSD1, which has been implicated in SCLC linage specification [7] and recognition by the immune system [8]. However, the heterogeneous drug responses in SCLC cells, arising from both intrinsic and acquired resistance to multiple LSD1 inhibitors, limit the potential of this strategy until biomarkers are identified to predict drug response [9]. Furthermore, recent studies have shown that chemotherapy-resistant SCLC cells exhibit elevated MYC expression and display distinct metabolic vulnerabilities. Thus, in murine models harboring MYC-driven tumors, targeted arginine depletion using pegylated arginine deiminase (ADI-PEG 20) significantly inhibited tumor progression and improved survival outcomes [10]. Inspired by the notion that epigenetic mechanisms, rather than druggable genetic driver,s determine adaptation to their microenvironment and response to therapy [11], epigenetic drugs like romidepsin or panobinostat provided a compelling rationale for treatment but ultimately failed to demonstrate a significant benefit for patients [12, 13]. In the light of these failures to translate preclinical concepts into improved disease control, SCLC remains regarded as a “graveyard of drug development” [14].

An underexploited concept in cancer therapy involves the use of redox-targeting drugs. The rationale behind this approach is that cancer cells are more vulnerable to therapy-induced reactive oxygen species (ROS) stress than non-cancerous cells, thereby creating an opportunity for selective drug targeting. We have recently identified a set of 15 genes, called “**A**nti-oxidant **C**apacity **B**iomarkers” (ACB) which accurately predict the sensitivity of cell lines across various cancer types to redox-targeting drugs such as ferroptosis inducers (GPX4 inhibitors) or thioredoxin reductase 1 (TXNRD1) inhibitors [15]. Importantly, only defined patient groups in each cancer entity express a favorable ACB profile (low-ACB expression), emphasizing the necessity of patient stratification for redox-targeting therapies.

In this study, we show that pathways protecting against ROS stress are underactive in SCLC cells. Upon treatment with a novel TXNRD1 inhibitor, these cells rapidly deplete their ROS-scavenging capacity, regardless of molecular subtype or cisplatin resistance status. Notably, unlike non-cancerous cells, SCLC cells are unable to compensate for TXNRD1 inhibition via the NRF2 stress response pathway. Our investigation into the biochemical and molecular basis of sustained drug sensitivity revealed a complex mechanism involving both epigenetic and transcriptional regulation. Leveraging this reduced ability to manage and adapt to oxidative stress, we conducted proof-of-concept studies in mouse models, demonstrating that TXNRD1 inhibition robustly maintained first-line therapy efficacy. Additionally, selective activation of the NRF2 stress response pathway using Bardoxolon-Methyl (CDDO-Me) allowed to increase the therapeutic dose of the TXNRD1 inhibitor, which improved tumor control without increasing toxicity in healthy tissues. These results highlight the therapeutic promise of TXNRD1 inhibitors for cancers like SCLC, characterized by diminished ROS-scavenging capacity and robust suppression of adaptive resistance mechanisms.

## Results

### SCLC cells are hypersensitive to TXNRD1 inhibition, independent of their resistance-status to cisplatin

We previously identified a set of 15 antioxidant capacity biomarkers (ACB), which accurately predict sensitivity to redox-targeting drugs like inhibitors of TXNRD1 and GPX4 [15]. While non-small cell lung cancer (NSCLC) cell lines demonstrate a wide range of ACB expression, we found that in SCLC, the majority of cell lines and PDX models express low levels of ACBs, indicative of high sensitivity to redox-targeting drugs (Fig. 1A, left panel, Fig. 1B, Supplementary Fig. 1A).

**Figure 1:**
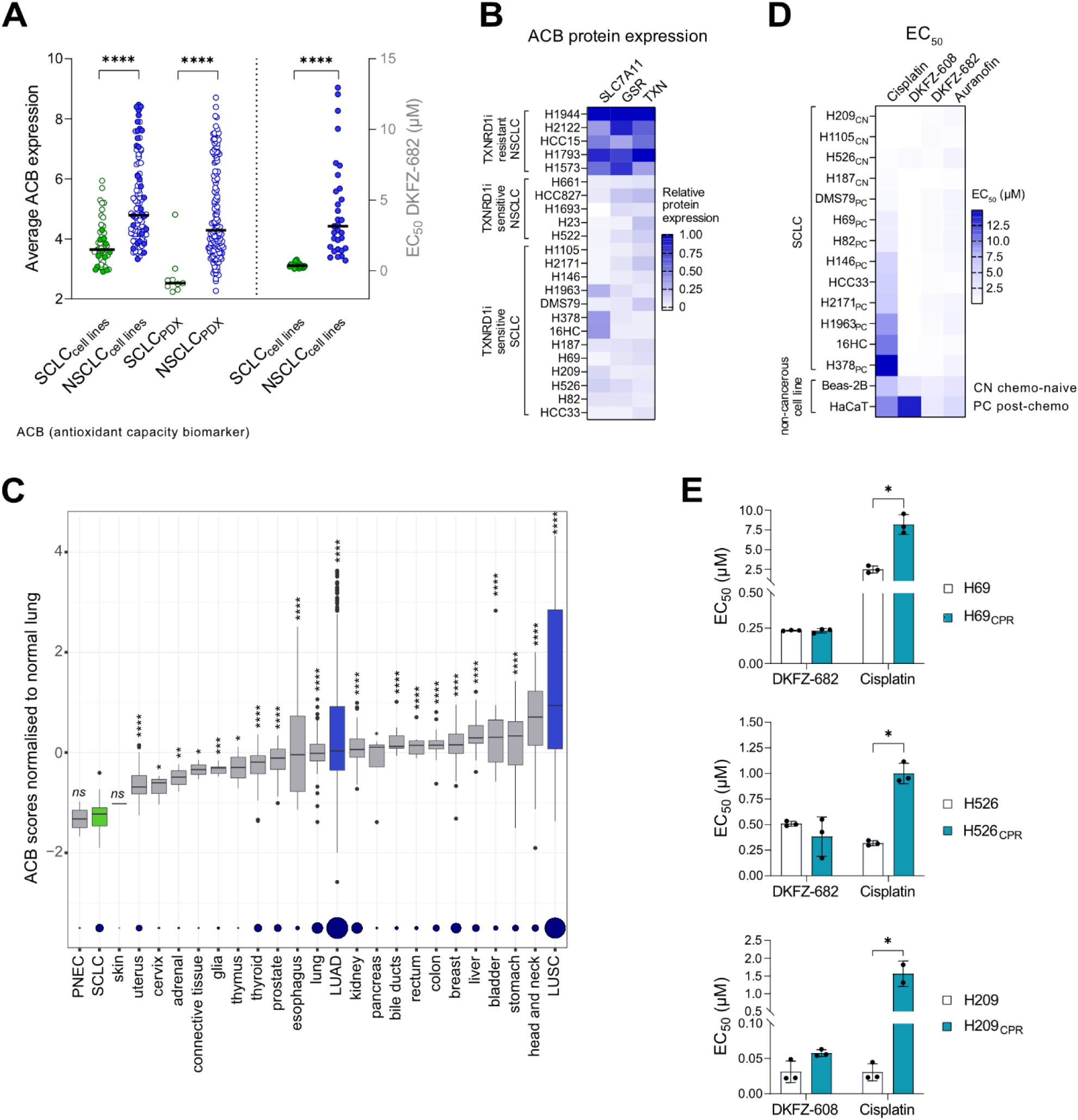
SCLCs have a low expression of ACBs and are very sensitive to TXNRD1 inhibition, regardless of their resistance-status to cisplatin. **(A)** The expression of 15 ACB genes for each SCLC and NSCLC cell line was calculated as average and are based on CCLE dataset and PDX, obtained from Champions Oncology (left panel). Units were converted to log2(TPM+1). To determine EC_50_ values, selected SCLC and NSCLC cell lines were treated with a concentration series of DKFZ-682 for 24 h and the cell viability was quantified by the CellTiter-Glo (SCLC) and CellTiter-Blue (NSCLC) assay (right panel). Dots represent mean of EC_50_ data from independent experiments performed in triplicates (n=2-6; \*\*\*\**p* < 0.0001, two-tailed unpaired *t*-test). **(B)** The heatmap with the protein levels of SLC7A11, GSR and TXN, analyzed in total cell extracts by immunoblotting. Protein expression in H1944 was set to 1 for each gene and for each experiment. Relative data represent mean of independent experiments (n=2-6). **(C)** Relative expression of ACB genes in normal tissues, LUSC, LUAD and PNEC were compared to SCLC using the Wilcoxon test (*ns*, not significant, \**p* < 0.05, \*\**p* < 0.01, \*\*\**p* < 0.001, \*\*\*\**p* < 0.0001). Dot size indicates sample size, ranging from n=1 (skin) to n=542 (LUAD). Expression values are normalized to normal lung tissue. **(D, E)** Cells were treated with a concentration series of cisplatin (72 h), DKFZ-608 (72 h), DKFZ-682 (24 h) and auranofin (24 h) and the cell viability was quantified by the CellTiter-Glo or CellTiter-Blue assay (H209/H209_CPR_ **(E)**). Data are presented as mean of EC_50_ values ± SD of independent experiments (n=2-4, \**p* < 0.5, two-tailed unpaired *t*-test) each performed in triplicates. **(D)** CN (chemo-naïve) SCLC cell lines are derived from chemo-naïve patients, PC (post-chemo) SCLC cell lines are derived from patients after cisplatin plus etoposide therapy. The therapy status of SCLC cells (HCC33 and 16HC) lacking labeling with “CN” or “PC” is uncertain. **(E)** H69_CPR_, H526_CPR_, H209_CPR_ are cisplatin-resistant sublines of NCI-H69, NCI-H526, and NCI-H209, respectively. Resistance to the drug in H69_CPR_ and H526_CPR_ was developed by gradually increasing the concentration of cisplatin in the growth medium once a week over a period of six months. H209_CPR_ was isolated from relapsed xenograft tumors in mice after cisplatin therapy.

Interestingly, the ACB status of SCLC is largely independent of the neuroendocrine subtype, with the exception of samples derived from PDX models (Supplementary Fig. 1B). Moreover we observed that the ACB levels are significantly lower in SCLC tumors as compared to normal tissues (Fig. 1C), suggesting that a selective tumor response to TXNRD1 inhibitors could be achieved in SCLC patients. Of note, pulmonary neuroendocrine cells (PNECs), which are the cells of origin of SCLC (reviewed in [16]), demonstrate similarly low-ACB scores, indicating that the ACB status of SCLC cells is inherited rather than selected during carcinogenesis.

To test whether this low-ACB status translates into high sensitivity to redox-targeting drugs, we assessed the activity of two TXNRD1 inhibitors, DKFZ-682 and DKFZ-608, in a panel of SCLC cell lines (n=14) that capture ACB expression levels present in 82% of this cancer entity (Fig.1A). The two molecules are equipotent enzymatic inhibitors of TXNRD1, show high correlation of cellular TXNRD1 inhibition and cytotoxicity and are, in contrast to auranofin, predominantly acting in the cytoplasm with limited mitochondrial ROS induction [15, 17] (Supplementary Fig. 1C-E). Our results show that unlike NSCLC, SCLC cells consistently exhibit sensitivity to TXNRD1 inhibition (Fig. 1A, right panel, Supplementary Fig. 1F), in line with the notion that low ACB levels translate into high activity of redox-targeting drugs in cancer. Furthermore, it highlights the potential of ACBs as a highly predictive biomarker for patient stratification, offering guidance on the use of oxidative stress induction as a therapeutic strategy.

The homogenous activity profile of our TXNRD1 inhibitors is especially notable compared to 1S3R-RSL3 (RSL3), which exhibits a more variable response pattern, independent of the cell’s ASCL1/REST or NEUROD1/REST status (Supplementary Fig. 2A, B). Overall, these findings suggest that TXNRD1 inhibition holds strong potential for SCLC therapy, given its consistent activity profile and independence from the neuroendocrine (NE) status of target cells.

In order to assess the efficacy of TXNRD1 inhibition in the context of platinum-based therapy, we assembled a panel of cells derived from both chemotherapy relapsed patients (PC, post chemo) and treatment-naïve patients (CN, chemo naïve) with varying resistance or sensitivity to cisplatin. PC-cells exhibit reduced expression levels of SLFN11 (Supplementary Fig. 2C), a previously described biomarker for chemotherapy resistance [18]. As such, these cells are approximately 20-fold more resistant to cisplatin compared to CN-cells (Fig. 1D, Supplementary Fig. 2D). Moreover, the cell line panel accurately reflects on the narrow therapeutic window of cisplatin following relapse, as we observe a ratio of only 1.7 of the average sensitivity of PC-SCLC compared to HaCaT and Beas-2B, which we used to represent non-cancerous cells. In contrast to cisplatin, TXNRD1 inhibitors DKFZ-682, DKFZ-608 and auranofin demonstrated equal efficiency in PC and CN cells, showing that TXNRD1 inhibitors are not affected by mechanisms causing cisplatin resistance (Fig. 1D). A comparison to non-cancerous cells revealed a therapeutic window of up to 50-fold (Supplementary Fig. 2D). The consistently high toxicity of TXNRD1 inhibitors across the entire SCLC cell line panel suggests an opportunity in overcoming resistance induced by first line therapy.

Therefore, we examined the effect of DKFZ-682 and DKFZ-608 on cells that survived cisplatin treatment (CPR, cisplatin-resistant), either selected in cell culture or derived from relapsed xenograft tumors in mice. In both cases, the cytotoxic efficacy of TXNRD1 inhibition remained unaffected (H69_CPR_, H526_CPR_) or only marginally decreased (H209_CPR_), while significant resistance to cisplatin was observed (Fig. 1E, Supplementary Fig. 3A, B). In addition, we tested primary circulating tumor cells (CTC), which were collected from relapsed SCLC patients [19]. Consistent with cell lines, CTCs exhibited a heterogeneous response to cisplatin with an inverse correlation of SLFN11 expression and drug sensitivity (r=-0.62) (Supplementary Fig. 2C) and a homogeneously high response to TXNRD1-inhibitor treatment, with DKFZ-608 being more efficient than auranofin (Supplementary Fig. 3C).

A recently published single cell data set [20] analyzing paired samples of CTCs from a naïve and relapsed patient revealed that ACB scores are only marginally increased in the cell population surviving cisplatin therapy (Supplementary Fig. 3D). Additionally, ACB levels do not segregate according to epithelial-mesenchymal transition (EMT) scores (Supplementary Fig. 3E), which have previously been shown to correlate with therapy resistance [21]. Our results demonstrate that SCLC cells are highly sensitive to TXNRD1 inhibitors and that cisplatin-resistant SCLC cells maintain this sensitivity, suggesting that TXNRD1 inhibition is a promising strategy for SCLC therapy.

Of note, the mechanism of cell death within the SCLC cell panel appears heterogeneous, despite the homogeneous toxicity induced by TXNRD1 inhibition. In HCC33 and H82 cells, TXNRD1 inhibition triggers an iron-dependent form of cell death, distinct from ferroptosis, as it can be rescued by the iron chelator desferrioxamine (DFO), but not by ferrostatin-1, which inhibits lipid peroxidation (Supplementary Fig. 4A). In H1105, in contrast to the afore mentioned cell lines, DKFZ-682 induces only a moderate increase of hydrogen peroxide (H_2_O_2_) and hydroxyl radical (HO·) (Supplementary Fig. 4B, C). Here, toxicity cannot not be rescued by DFO but by pretreatment with olaparib, suggesting the involvement of PARP-dependent apoptosis (Supplementary Fig. 4D; more experimental details are described in Supplementary section 1 and Supplementary Fig. 4E-G).

### Targeting TXNRD1 maintains efficacy of first-line therapy *in vivo*

SCLC patients typically show high initial response rates to cisplatin-based chemotherapy, followed by relapse shortly after completing first-line therapy. Given the significant efficacy of TXNRD1 inhibition observed in both naïve and cisplatin-resistant cancer cells, we hypothesized that our TXNRD1 inhibitors could be effective as maintenance therapy following first-line therapy with cisplatin/etoposide. To test this hypothesis, we used our preclinical xenograft model with the human SCLC cell line H209, which was derived from an untreated patient and asked whether DKFZ-608 could prevent tumor recurrence following complete remission after standard chemotherapy with cisplatin/etoposide. The experimental setup and the therapeutic regimen closely mirror those used in most previously published SCLC xenograft studies, with the key difference that we extended monitoring of tumor-free survival (time to recurrence) for an additional 4 months beyond the end of treatment to detect delayed recurrence (Fig. 2A-C).

**Figure 2:**
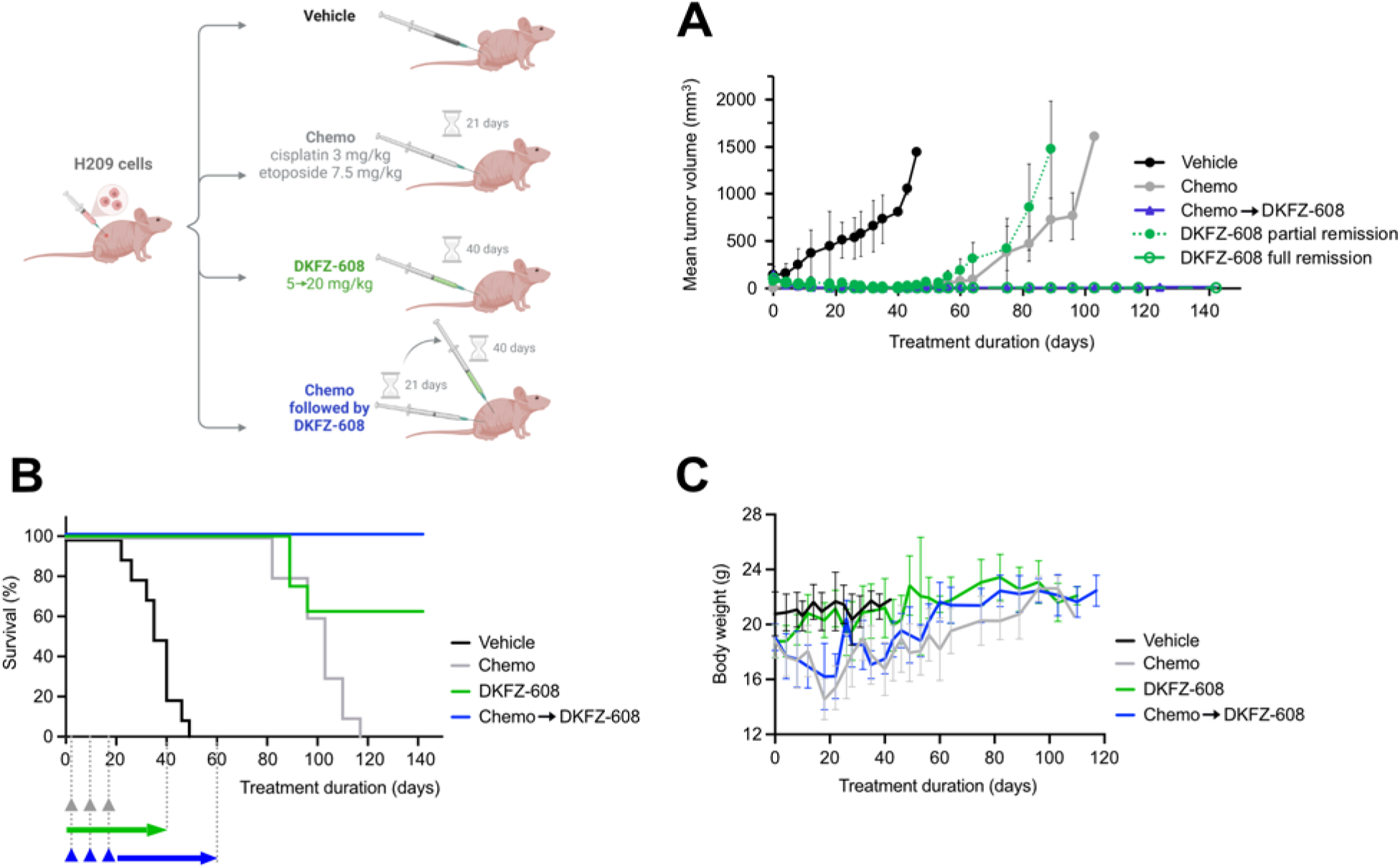
Targeting of TXNRD1 is efficient as a maintenance therapy for chemotherapy-resistant SCLC tumors *in vivo*. Human SCLC cell line H209 was transplanted subcutaneously into nude mice. Tumor-bearing mice were distributed into groups (n=10) and treated with cisplatin/etoposide (chemo) for 3 weeks or DKFZ-608 for 40 days. One group started on the chemo regimen for 3 weeks followed by DKFZ-608 for 40 days. The survival **(A),** tumor size **(B)** and body weight **(C)** were monitored during the course of the experiment. The mean values ± SD of the tumor size and body weight are shown. In **(B)**, the group receiving DKFZ-608 mono-therapy (green) is presented as two separate lines in order to allow a better presentation of animals reaching full remission.

As expected, the tumors in the control group (vehicle only, black line, Fig. 2A), showed rapid growth, necessitating the sacrifice of mice within 37 ± 9 days. In contrast, the second group that received 3 cycles of cisplatin (3mg/kg, interperitonally, i.p.) and etoposide (7.5 mg/kg, i.p.), responded with rapid tumor shrinkage (gray line, Fig. 2A). After 20 days, tumors were undetectable in 9 out of 10 animals, leading to the cessation of treatment. However, 18 days later, measurable tumors reappeared in all animals with the first animal reaching maximum tumor size of 1500 mm^3^ within 60 days after treatment discontinuation. The mean survival time, calculated from the start of the experiment, was 100 ± 11 days (Fig. 2B).

The third group received DKFZ-608 (15 mg/kg daily, i.p.) immediately following chemotherapy as a maintenance treatment for 40 days (blue line, Fig. 2A, B). Notably, no tumor recurrence was observed until 96 days after the end of chemotherapy or 56 days after discontinuation of DKFZ-608. Thereafter, we observed a faint regrowth of the tumor in a single animal. The remaining nine mice were tumor free until day 142, at which point the experiment was concluded.

The fourth group, receiving monotherapy of DKFZ-608 (15 mg/kg) also showed tumor shrinkage (green line, Fig. 2A). By the end of the treatment (day 43), 6 animals were tumor-free, while the remaining 4 showed approximately 70% tumor shrinkage. In these four animals, tumor growth resumed after discontinuation of therapy, while no recurrence was observed in the six tumor-free animals during the 142-day experimental period.

The preclinical data presented here clearly demonstrate that the TXNRD1 inhibitor DKFZ-608 is a promising drug candidate for the treatment of SCLC, effectively targeting both first-line sensitive and resistant SCLC cells. It is capable of completely suppressing tumor relapse for several months at a sub-toxic dose. To the best of our knowledge, this is the first instance of such a robust remission being achieved in an SCLC mouse model.

### Reduced ROS-scavenging capacity mediates vulnerability to redox-targeting drugs in SCLC

The activity of NRF2 is widely recognized as a key determinant of resistance to many cancer drugs (reviewed in [22]). The hypersensitivity of SCLC cells to TXNRD1 inhibition raises the question of whether NRF2-mediated ROS defense is compromised in these cells. SCLC and NSCLC cells have similar NRF2 transcript expression levels, but significantly different NQO1 mRNA levels (Supplementary Fig. 5A, B). Considering NQO1 mRNA levels as a proxy for NRF2 activity [23] indicates that SCLC cells exhibit on average lower NRF2 activity than NSCLC cells. This finding aligns with the low NRF2 protein levels observed in our SCLC cell line panel (Supplementary Fig. 5C), suggesting that low steady-state levels of redox-scavenging is determined by low NRF2 activity.

Interestingly, that SCLC cells, despite their low ACB expression, show similar or even lower steady-state levels of hydrogen peroxide (H_2_O_2_) and oxygen radicals (OH· and O_2_·^-^), as compared to cells with high ROS-scavenging capacity such as high-ACB NSCLC cells (Supplementary Fig. 6A, B). However, in response to TXNRD1 inhibition, ROS levels increase dramatically in most SCLC cells, while remaining unchanged in high-ACB NSCLC cells, supporting the hypothesis that SCLC cells have a reduced ROS-scavenging capacity (Supplementary Fig. 6C, D). We also observed that SCLC cells have higher intracellular levels of reactive nitrogen species (RNS), such as nitric oxide (NO) and peroxynitrite (ONOO^-^), compared to high-ACB cells (Supplementary Fig. 6E-G), which may be necessary to support their proliferation [15].

We next explored whether the NRF2 system in SCLC is capable of responding to inhibition of KEAP1, the E3 ligase responsible for regulating NRF2 protein levels. As anticipated, treatment with CDDO-Me and other KEAP1 inhibitors led to an increase in NRF2 protein levels in both SCLC cells and non-cancerous Beas-2B and HaCaT cells (Fig.3A, Supplementary Fig. 7A-C). Among SCLC cell lines, H82 and H526 exhibited the highest NRF2 protein levels upon KEAP1 inhibition along with marked upregulation of the NRF2 target genes NQO1 and AKR1C3, indicating that the NRF2 regulon is functional in those two cell lines (Fig. 3B, C). However in contrast to non-cancerous cells, the majority of ACBs and other genes of the redox regulatory system remained at low expression levels, resulting in only marginal activation of pathways involved in ROS defense (Fig. 3D, E). Consistently, SCLC cells failed to increase their ROS-scavenging capacity to adapt to TXNRD1 inhibition (Fig. 3F, Supplementary Fig. 7D). We specifically examined the role of glutathione (GSH) and nicotinamide adenine dinucleotide phosphate (NADPH), which are essential for maintaining ROS homeostasis. In Beas-2B cells, the depletion of GSH by inhibition of glutathione synthesis (buthionine sulphoximine, BSO) induces cell death, which is rescued by CDDO-Me, although GSH levels remain unchanged (Supplementary Fig. 8A, B). Notably, the protective effect of CDDO-Me is lost when NADPH regeneration via G6PD is also inhibited with G6PDi-1 (Supplementary Fig. 8B) [24]. Similarly, under conditions of DKFZ-682-induced ROS stress, the protective effect of CDDO-Me is negated only when cells are treated with both BSO and G6PDi-1 (Supplementary Fig. 8C), suggesting that NADPH-dependent pathways can partially compensate when GSH-mediated redox homeostasis is exhausted. In drug-sensitive SCLC, GSH levels appear elevated compared to more resistant non-cancerous cells (Supplementary Fig. 8D), suggesting that both steady-state and induced GSH levels may be insufficient to counteract drug-induced ROS-stress. The capacity to scavenge ROS seems to be mainly limited by NADPH levels, which are significantly reduced in SCLC cells, in line with their low expression of G6PD (Supplementary Fig. 8E, F).

**Figure 3:**
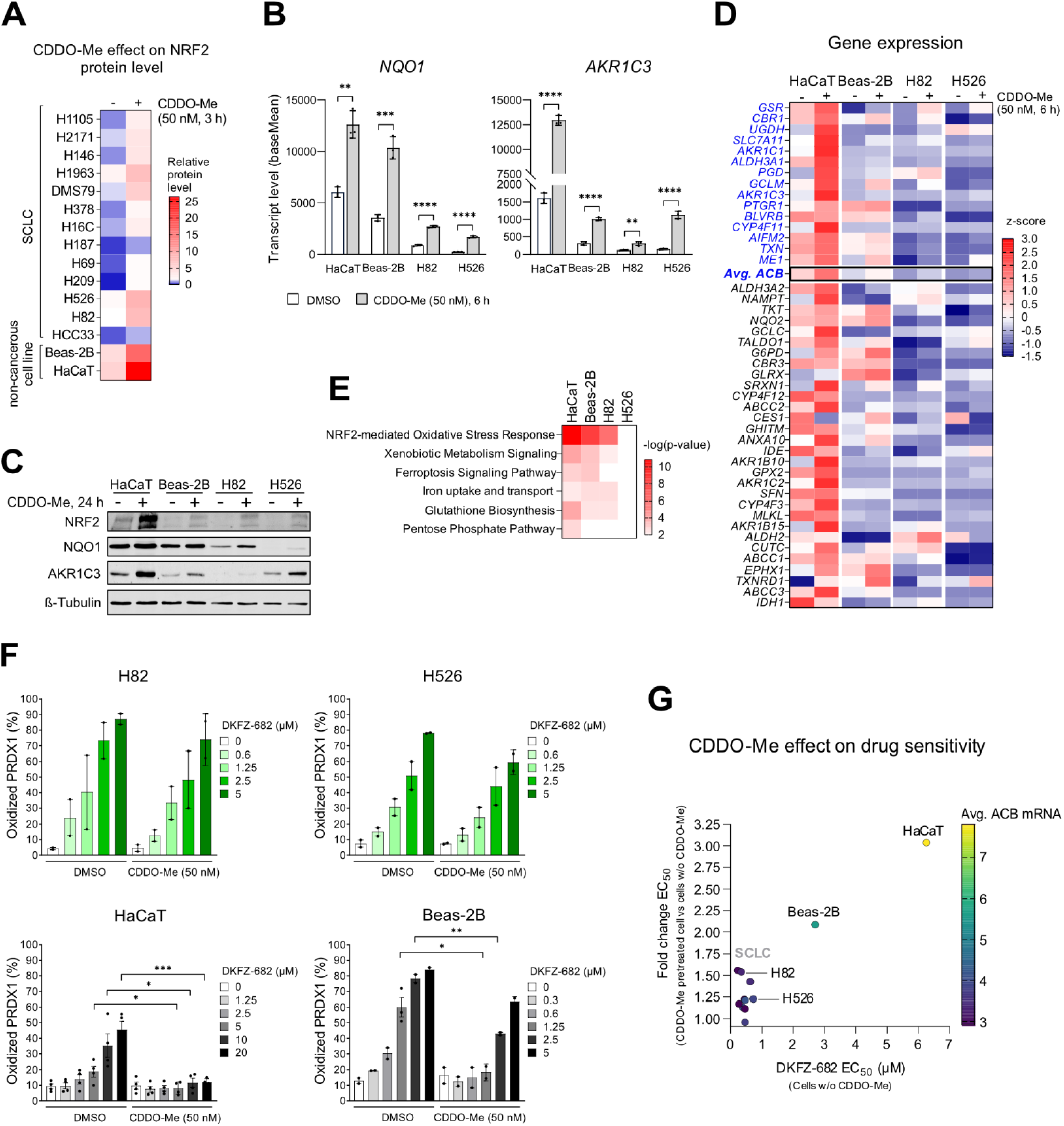
The activation of NRF2 pathway by CDDO-Me protects non-cancerous cells but not SCLCs against the cytotoxic effect of TXNRD1 inhibitors. **(A)** The NRF2 protein level in total cell extracts was analyzed in SCLC and non-cancerous cell lines upon treatment with DMSO (control) or CDDO-Me by immunoblotting. NRF2 expression in DMSO treated H82 cells was set to 1 for each experiment. Relative data represent mean of independent experiments (n=2-4). **(B)** The induction of *NQO1* and *AKR1C3* transcripts upon CDDO-Me treatment in different cell lines was determined by expression profiling (n=3, mean ± SD; \*\**p* < 0.01, \*\*\*\**p* < 0.0001, two-tailed unpaired *t-*test). **(C)** The induction of NQO1 and AKR1C3 proteins upon CDDO-Me (50 nM) treatment was determined by immunoblotting (representative of 3 independent experiments). **(D)** The top 45 genes highly correlated with resistance to TXNRD1 inhibition [15] were analyzed by expression profiling of two non-cancerous and two SCLC cell lines treated with CDDO-Me compared to a DMSO control (n=3). ACBs are labelled in blue. **(E)** The most up-regulated pathways upon CDDO-Me treatment according to a pathway enrichment analysis based on the expression profiling data. **(F)** Oxidized and reduced levels of PRDX1 protein were analysed by immunoblotting in cells treated first with CDDO-Me (50 nM) for 24 h and then with the indicated concentrations of DKFZ-682 for 3 h. Bar diagrams summarize the quantitative results from independent experiments (n=2-4, mean ± SD; \**p* < 0.05, \*\**p* < 0.01, \*\*\**p* < 0.001, two-tailed unpaired *t-*test). **(G)** The cells were treated with CDDO-Me (50 nM) or DMSO (control). After 24 h, the cells were treated with a concentration series of DKFZ-682 for 24 h and the cell viability was measured by the CellTiter-Glo assay. The fold change of EC_50_ for DKFZ-682 upon CDDO-Me compared to cells without CDDO-Me pretreatment was calculated (n=2-4).

Most significantly, the differential response to KEAP1 inhibition observed in SCLC compared to non-cancerous cells effectively broadens the cellular therapeutic window of TXNRD1 inhibition. SCLC cells remain largely sensitive despite NRF2 induction, whereas non-cancerous cells exhibit increased resistance (Fig. 3G). The absence of resistance induction upon KEAP1 inhibition in SCLC cells was unexpected, given that ROS-induced inactivation or genetic loss of KEAP1 is a common mechanism of resistance to redox-targeting drugs in many cancer types [22]. One possible explanation is the constitutive nuclear activity of BACH1, a transcriptional repressor of a subset of NRF2 target genes [25], or the lack of a positive cofactor like MAFG [26]. However, our detailed analysis revealed that neither BACH1 nor MAFG is predominantly responsible for the lack of adaptation to drug induced ROS stress (experimental details in Supplementary Section 2, Supplementary Fig. 9, 10).

An alternative explanation for the lack of NRF2-response could be the epigenetic silencing of redox genes. Using public data sets of CpG clusters [27], we observed that several ACB promoters, in particular ME1, PTGR1, PGD and ALDH3A1 show a higher degree of methylation in SCLC and low-ACB NSCLC cells compared to high-ACB NSCLC cells. This suggests that epigenetic silencing could contribute to the failure to adapt to oxidative stress (Fig. 4A). To avoid loss of resolution caused by CpG clustering, we performed methylation analysis on 3 SCLC and 3 NSCLC cell lines, quantifying individual CpG loci. Our results indicate that TXN1, a member of the ACB gene set, may also be subject to epigenetic silencing. We identified a high degree of DNA methylation at positions −995 and −1418 (CpG1 and CpG2 respectively) in the distal promoter region of *TXN1*, both in cell lines and patient tumors. This region has previously been shown to contain an oxidative stress response element [28] (Fig. 4B, C). Consistent with this, TXN1 protein levels are strongly reduced in SCLC compared to NSCLC tumors, which exert lower *TXN1* promoter methylation (Fig. 4D).

**Figure 4:**
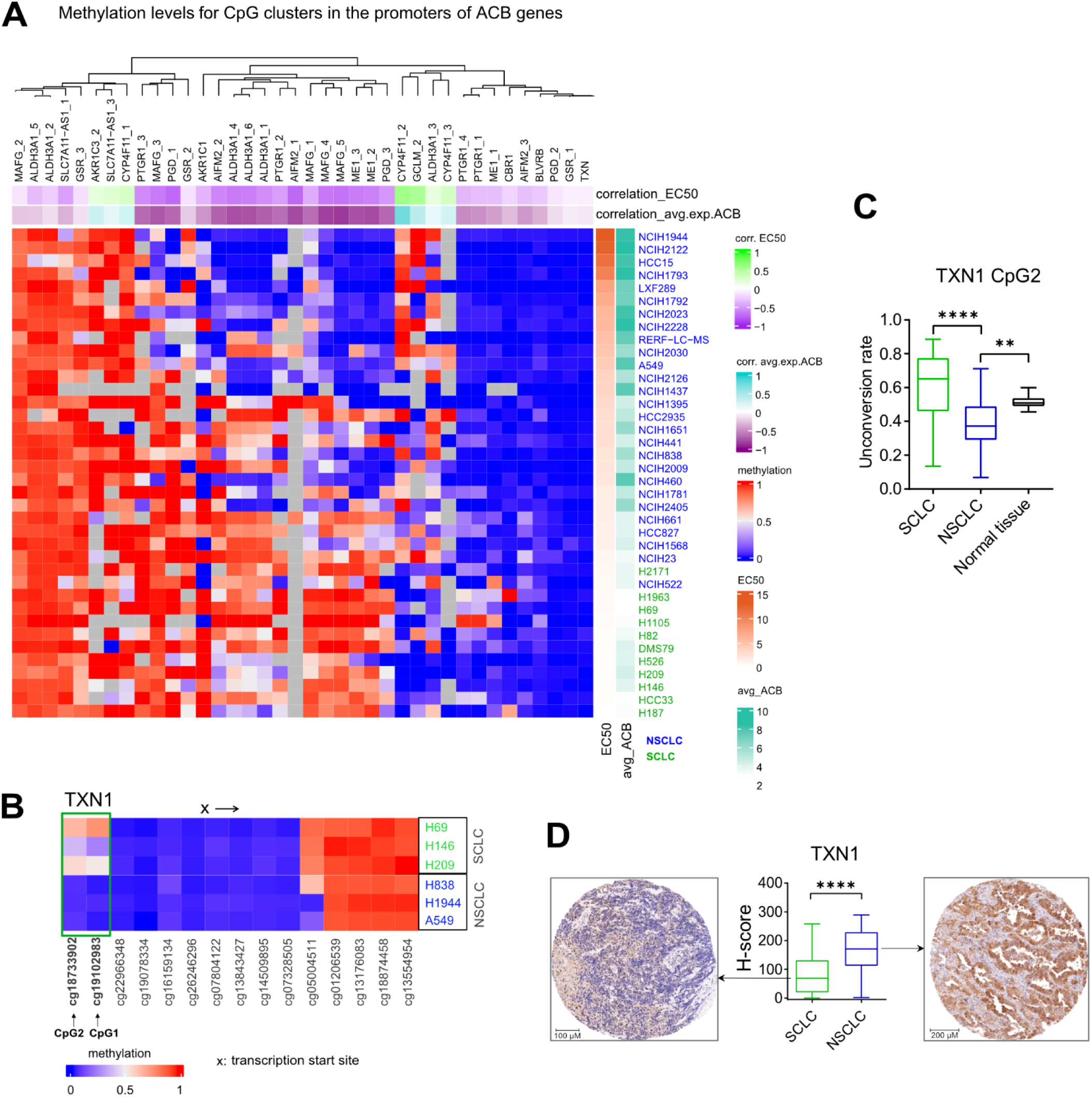
Increased promoter methylation of ACB genes in SCLCs correlates with the sensitivity to TXNRD1 inhibitor. **(A)** Methylation values for CpG clusters within promoters of MAFG and ACB gene set for NSCLC and SCLC cell lines profiled in DepMap project (CCLE) that have expression and promoter methylation data. Row annotation: 1st line – DKFZ-682 EC_50_ per cell line (rows are sorted based on these values); 2nd line – average expression of genes from ACB gene set per cell line. Column annotation: 1st line – correlation between vector of cell lines’ methylation values and vector of cell lines’ DKFZ-682 EC_50_ values per each CpG cluster; 2nd line – correlation between vector of cell lines’ methylation values and vector of cell lines’ average expression of genes from ACB gene set per each CpG cluster. **(B)** The methylation status of genomic DNA was analyzed using the Infinium MethylationEPIC BeadChips (Illumina). The heatmap depicts beta values with 0 meaning no methylation and 1 indicating high methylation status. The probes are indicated by Illumina IDs and sorted according to their localization in the *TXN1* gene. CpG1 and CpG2 are indicated by their distance to the transcription start site. **(C)** The methylation levels of CpG2 in *TXN1* promoter were analyzed in samples from SCLC (n=40) and NSCLC (n=46) patients using bisulfite sequencing PCR. Normal tissue samples originating from NSCLC patients (n=10) were used as a control. Box plots depict unconversion ratios with 0 standing for no methylation and 1 indicating high methylation status (\*\**p* < 0.01, \*\*\*\**p* < 0.0001, two-tailed unpaired *t*-test). **(D)** The TXN1 levels of SCLC (n=72) and NSCLC (n=88) specimen were determined based on the digital evaluation of TXN1 immunohistochemical staining using tissue microarrays. Based on the staining intensities H-scores were calculated (\*\*\*\**p* < 0.0001, two-tailed unpaired *t*-test). Tumor specimen representative for the median H-score of each group are depicted.

To test whether hypermethylation was indeed involved in the lack of NRF2 responsiveness of ACB promoters, we pre-treated H82 and H69 cells with DNA methyl transferase inhibitors decitabine (DAC) or 5-aza-2′-deoxycytidine (AZA), followed by CDDO-Me treatment. Contrary to our expectation, DNA-demethylation did not lead to more efficient drug resistance by NRF2 induction (Supplementary Fig. 11A), in line with inconsistent induction of hypermethylated ACB-promotors (Supplementary Fig. 11B). This indicates that, despite the high correlation of promoter methylation with ACB expression and drug sensitivity, epigenetic silencing is not a primary course for the lack of ACB induction by NRF2 and the inability of SCLC to adapt to drug-induced ROS stress.

In summary, SCLC cells suppress ROS defense enzymes through overlapping layers of epigenetic and transcriptional mechanisms. Consequently, SCLC cells exhibit a reduced ROS-scavenging capacity and, unlike non-cancerous cells, are unable to adapt to drug-induced ROS stress via NRF2-mediated mechanisms.

### Widening the therapeutic window for redox-targeting drugs

Given that SCLC cells are unable to adapt to redox-targeting drugs, such as TXNRD1 inhibitors, we hypothesized that activating the NRF2 pathway would enhance antioxidant defenses specifically in non-cancerous cells, thereby further widening the therapeutic window. To test this hypothesis *in vivo*, we utilized the hydroxy-substituted derivative DKFZ-682, which exhibits slightly lower potency against SCLC and reduced selectivity towards non-cancerous cells, compared to DKFZ-608 (Supplementary Fig. 2D). Consequently, DKFZ-682 has a lower maximum tolerated dose (MTD), limiting its efficacy in eliminating cancer cells.

Upon treatment of mice with the KEAP1 inhibitor CDDO-Me at a concentration previously established for the i.p. route [29] (Fig. 5A), we observed the induction of NRF2 proxies, such as NQO1, GSR and GCLM in the liver, kidney and lung (Fig. 5B). On a physiological level, we observed a reduction of stress markers, including blood urea nitrogen (BUN), alanine aminotransferase (ALT), and aspartate aminotransferase (AST) (indicators of kidney and liver damage), induced by DKFZ-682 (Fig. 5C), supporting the hypothesis of organ protection.

**Figure 5:**
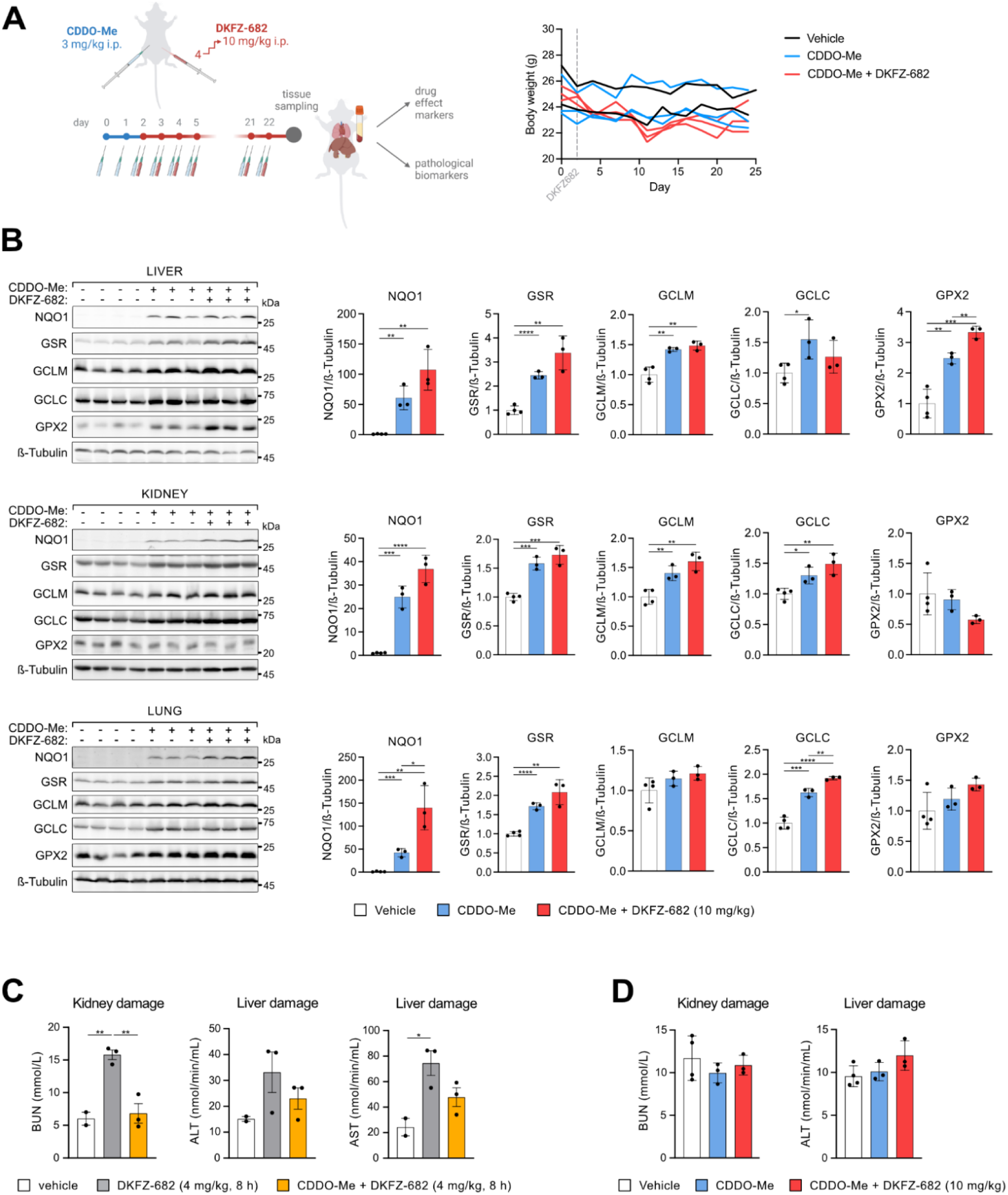
CDDO-Me widens the therapeutic window for a TXNRD1-targeting inhibitor *in vivo*. **(A)** After CDDO-Me pre-treatment, the NSG mice were daily injected intraperitoneally with DKFZ-682, gradually increasing the dose from 4 mg/kg to 10 mg/kg that was then maintained for up to 3 weeks. The body weight of individual animals was monitored during the course of the experiment. **(B)** The induction of drug effect markers of CDDO-Me in mouse tissues was investigated by immunoblotting (n=3-4). The bar graphs show quantification of signal intensities (mean ± SD, \**p* < 0.05, \*\**p* < 0.01, \*\*\**p* < 0.001, \*\*\*\**p* < 0.0001, two-tailed unpaired *t*-test). **(C)** The organ damage markers were determined in the blood plasma of mice after a single injection of 4 mg/kg DKFZ-682 with or without CDDO-Me pre-treatment for 48 h (n=2-3, mean ± SD, \**p* < 0.05, \*\**p* < 0.01, two-tailed unpaired *t*-test). As markers of kidney and liver damage, blood urea nitrogen (BUN) and alanine aminotransferase (ALT) as well as aspartate aminotransferase (AST) were measured, respectively. **(D)** The organ damage markers were determined in the blood plasma after the 3-week treatment as presented in **A** (n=3-4). As markers of kidney and liver damage, BUN and ALT were measured, respectively.

Consistent with this, we were able to increase the therapeutic dose by 2.5-fold without causing disproportionate stress or weight loss (Fig.5A, right panel). Remarkably, stress markers remained suppressed over a period of 3 weeks, indicating that NRF2-mediated organ protection can be maintained long-term without desensitization (Fig. 5D).

We next compared tumor growth reduction in unprotected (MTD 4 mg/kg DKFZ-682) with CDDO-Me-protected mice (MTD 10 mg/kg DKFZ-682) (Fig. 6A). To monitor drug efficacy more precisely, we administered DKFZ-682 as monotherapy rather than maintenance therapy, allowing us to eliminate confounding effects from first-line therapy. In addition, we selected a tumor model based on H526, which exhibits some resistance to TXNRD1 inhibition (Supplementary Fig. 2D), to prevent complete suppression of tumor growth at the MTD of 4 mg/kg in unprotected mice.

**Figure 6:**
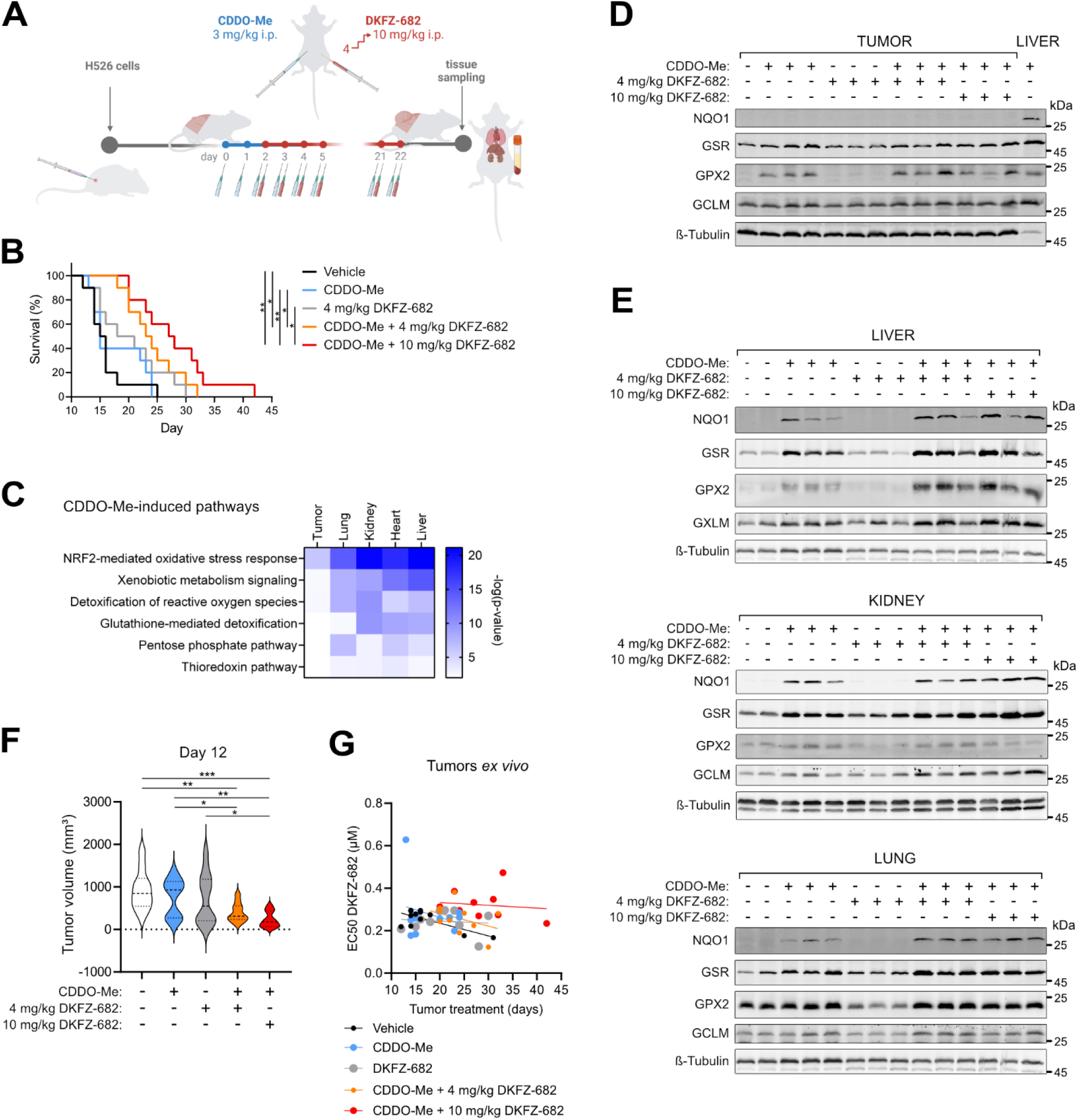
A selective activation of NRF2 pathway by CDDO-Me in normal tissues expands the therapeutic potential of TXNRD1 inhibitors *in vivo*. **(A)** As a tumor model, NSG mice were engrafted subcutaneously with the human SCLC cell line H526. After CDDO-Me pre-treatment, the NSG mice were daily injected intraperitoneally with DKFZ-682, gradually increasing the dose from 4 mg/kg to 10 mg/kg that was then maintained for up to 3 weeks. The tumor size was monitored daily for reaching the humane end-points. **(B)** Mice were randomly distributed into groups (n=10) that were daily injected with vehicle, CDDO-Me (3 mg/kg), a low dose of DKFZ-682 (dose escalation from 1 mg/kg to 4 mg/kg), CDDO-Me (3 mg/kg) and a low dose of DKFZ-682 (4 mg/kg), CDDO-Me (3 mg/kg) and a high dose of DKFZ-682 (dose escalation from 4 mg/kg to 10 mg/kg). The plot shows the survival of individual groups (\**p* < 0.05, \*\**p* < 0.01, log-rank (Mantel-Cox) test). **(C)** The pathways up-regulated by CDDO-Me were analyzed in mouse tissues and tumors performing RNAseq. Samples from tumor-bearing mice (H526 tumors) that were injected with CDDO-Me (3 mg/kg) for 4 days were compared to vehicle (n=5). **(D, E)** The induction of NRF2 target proteins in tumors **(D)** and mouse tissues **(E)** upon CDDO-Me and DKFZ-682 treatment was investigated by immunoblotting. Representative western blots are shown. **(F)** The tumor size measured 12 days after the daily mouse treatment was started (n=10, one-way ANOVA *p* = 0.0025; \**p* < 0.05, \*\**p* < 0.01, \*\*\**p* < 0.001 two-tailed unpaired *t*-test). **(G)** The sensitivity to DKFZ-682 was tested in explanted tumors after mouse treatment for the indicated time. Tumor cells were treated with a concentration series of DKFZ-682 for 24 h and cell viability was quantified using CellTiter-Glo. Each dot represents a tumor from one mouse measured in a triplicate.

Upon detection of palpable tumors, mice were randomized into groups receiving either vehicle or compounds daily for 3 consecutive weeks. The groups treated with CDDO-Me received injections for 2 days prior to the initiation of DKFZ-682 treatment (Fig. 6A). As anticipated, the antitumor response of DKFZ-682 at 4 mg/kg (Fig. 6B, gray line) was only moderate, resulting in a median survival of 20 days compared to 15.5 days in the vehicle-treated group (p=0.4) (Fig. 6B, black line). In contrast, the group treated with high-dose DKFZ-682 (10 mg/kg, Fig. 6B, red line) showed enhanced efficacy, with median survival increasing nearly 2-fold from 15 days in the CDDO-Me-only group (blue line) to 27 days in the CDDO-Me/high-dose DKFZ-682 group (p=0.001). These findings confirm our hypothesis that tolerance to an increased therapeutic dose can indeed translate into improved tumor-suppressive effects.

Pathway analysis of RNAseq data from organs and tumors demonstrated substantial induction of the NRF2 regulon in the lung, kidney, heart, and liver, in particular genes involved in redox homeostasis, but only a weak response in tumors (Fig. 6C). This differential response was not due to reduced efficacy of CDDO-Me in tumors since we detected substantial induction of the NRF2 proxy GPX2, at both protein (Fig. 6D) and RNA levels (Supplementary Fig. 12A).

Interestingly, mice that did not receive CDDO-Me protection during the 3 weeks of low-dose DKFZ-682 treatment failed to express higher levels of ROS response markers such as NQO1, GPX2, GCLM, or GSR in their organs (Fig. 6E), suggesting that sustained NRF2-mediated adaptation to drug-induced ROS stress does not occur without pharmacological KEAP1 inhibition.

Previous studies using mouse models of lung and prostate cancer have demonstrated that CDDO-Me exhibits antitumor effects and enhances the efficacy of combination therapies (reviewed in [30]). Contrary to these findings, our results indicated a positive interaction between CDDO-Me and the cytotoxic drug only during the early phase of the trial (Fig. 6F), however without a corresponding improvement in long term survival (Fig. 6B, orange versus gray line). Importantly, tumor cells isolated from CDDO-Me-treated mice did not show significant signs of acquired resistance to DKFZ-682 (Fig. 6G) and remained unresponsive to NRF2 induction when tested *ex vivo*. This suggests that CDDO-Me treatment did not promote the development of resistance in SCLC tumors (Supplementary Fig. 12B).

Taken together, our data provide evidence that selective activation of the KEAP1/NRF2 pathway in normal tissues minimizes dose-limiting toxicity and expands the therapeutic potential of TXNRD1 inhibitors. This approach paves the way for a paradigm shift in the treatment landscape of SCLC and in the use of ROS-inducing drugs in cancers with an impaired NRF2 pathway.

## Discussion

Despite decades of intense research and numerous clinical trials investigating targeted therapeutics, drug combinations, and immune checkpoint inhibitors, treatment options for SCLC patients remain limited. Maintenance therapy, however, holds significant promise for advancing SCLC treatment. Standard therapy with cisplatin/etoposide is highly effective, often resulting in near-complete remission, thus creating an ideal window for suppressing residual cancer cells. Unfortunately, none of the therapies tested so far have prevented tumor relapse. Successful maintenance therapy must target a critical vulnerability shared across all tumor cells present after first-line therapy and remain effective against emerging resistance mechanisms. The limited success of previous attempts suggests that the drug combinations have either failed to effectively target such a vulnerability in first-line-treated SCLC cells or were compromised by the same resistance mechanisms that blunted the efficacy of cisplatin/etoposide.

The TXNRD1 inhibitor DKFZ-608 exhibits a unique activity profile. In tumors derived from therapy-naïve cells, we observed sustained remission in mice. Remarkably, TXNRD1 inhibition appears fully compatible with first-line therapy, as it neither delayed the recovery of animals from chemotherapy-induced side effects nor caused cumulative toxicity, even after 40 consecutive daily doses (Fig. 2C). We identified several factors contributing to the remarkable efficiency of TXNRD1 inhibition on SCLC tumors. First, targeting redox homeostasis via TXNRD1 inhibition appears to hit an Achilles heel in SCLC, which is hallmarked by an inherently low steady-state capacity to scavenge drug-induced ROS stress. Additionally, SCLC cells are unable to upregulate their multimodal ROS-scavenging pathways to adjust to this stress (Fig. 3). Another, particularly important factor is that SCLC cells retain their low ROS buffer capacity and sensitivity to TXNRD1 inhibition even after acquiring resistance to chemotherapy. This is evidenced by the uniform potency of TXNRD1 inhibition across a panel of cell lines and circulating cancer cells, which exhibit heterogeneous responses to cisplatin (Fig. 1). We did not expect this, as cisplatin not only affects DNA replication but also induces ROS, thus contributing to its efficacy on tumor cells and systemic toxicity (reviewed by [31] and [32]). The events leading to cisplatin resistance in SCLC are still not fully understood, with conflicting reports on adaptive changes in redox systems (reviewed in [33]). Notably, the fact that patient-derived paired samples of pre- and post-therapy SCLC cells show comparable levels of ACBs encourages us to hypothesize that breaking of cisplatin-resistance can also be achieved in real-world SCLC maintenance therapy.

A third reason for the high potential of TXNRD1-targeting drugs is their efficacy, which is independent of the molecular subtype of SCLC. This contrasts with the heterogeneous toxicity of GPX4 inhibitors, which have previously shown resistance in certain NE subtypes [34]. In agreement with this earlier report, we observed up to a 250-fold difference in drug sensitivity to RSL3 in our SCLC cell line panel, highlighting the more homogeneous activity profile of TXNRD1 inhibitors (Supplementary Fig. 2A, B, D).

Why do SCLCs exhibit a low ROS-scavenging capacity and how is this achieved? Our data show that SCLC cells maintain low levels of basal ROS such as O_2_·^-^, HO· and H_2_O_2_ (Supplementary Fig. 6). One possible explanation for this phenomenon is that SCLC cells generate lower levels of O_2_^-^ through NADPH oxidases or mitochondrial respiration, which reduces the need for a high ROS-scavenging capacity to sustain the redox homeostasis required for rapid proliferation. This hypothesis is consistent with a recent report showing that SCLC cells have significantly fewer mitochondria, compared to NSCLC [35]. Reduced enzymatic ROS-scavenging could, at least under homeostatic conditions, be compensated by elevated levels of NO, which we detect in SCLC cells and which may act as a ROS scavenger by reacting with superoxide to form the non-radical, slow-reacting molecule peroxynitrite (ONOO^−^, reviewed in [36] and [37]). Additionally, SCLC cells may utilize hydropersulfide-based mechanisms, which have recently been shown to eliminate intracellular radicals [38]. Notably, among lung cancer cell lines, SCLC-derived cells express higher levels of cystathionine beta-synthase (CBS), the rate-limiting enzyme of the trans-sulfuration pathway, involved in the production of cysteine and hydrogen sulfide, which are essential for the formation and function of hydropersulfides (reviewed in [39]).

An alternative hypothesis is inspired by the finding that PNEC, the proposed cells-of-origin for NE SCLC, also exhibit a low-ACB profile (Fig. 1C). Oxygen sensing by PNECs relies on the precise regulation of H_2_O_2_, which is produced by an oxygen-consuming NADPH oxidase. H_2_O_2_ functions as a second messenger that modulates the activity of oxygen-sensitive K^+^ channels [40]. To accurately sense changes in H_2_O_2_ levels both under normoxic and hypoxic conditions, it is plausible that PNECs may limit their response to fluctuating H_2_O_2_ concentrations by avoiding the activation of adaptive ROS-scavenging pathways.

Currently, it is enigmatic how PNECs or SCLC cells establish a low ROS buffer status. Redox homeostasis is regulated by multiple transcription factors [41], with NRF2 playing a major role both in steady-state conditions and, more prominently, under ROS-induced stress. The fact that NRF2 mRNA levels are comparable to non-cancerous cells but NRF2-response genes, like NQO1 and the ACB gene set, are under-expressed suggests that SCLC cells have constitutively low NRF2 activity, most likely mediated by post-translational mechanisms like KEAP1-mediated degradation. Although we did not formally investigate these options, we also consider additional mechanisms that may modulate the activity of KEAP1 to contribute to low steady-state NRF2 activity in SCLC: Based on DEPMAP data, Sequestosome 1 (SQSTM1/p62), an inhibitor of KEAP1 [42], appears to be under-expressed in SCLC cells, while BRD4 [43], an activator of KEAP1 expression, is overexpressed in SCLC-derived cells.

The limited capacity to scavenge drug induced ROS observed under steady-state conditions is only one aspect contributing to the effectiveness of TXNRD1 inhibitors. An equally important factor in real-world therapy is adaptation to drug-induced ROS stress. Our data demonstrate that SCLCs are unable to adjust their ROS buffer capacity to drug-induced ROS challenge. We still have only a fragmented understanding of the underlying mechanisms. Our findings show that the hypermethylation of ACB promoter genes, while strongly correlated with ACB expression and drug sensitivity, is not a primary cause of impaired adaptation to ROS stress. We hypothesize that SCLC may lack a cofactor needed by NRF2 for efficient ACB induction. FOSL1 (FRA1), which is expressed at very low levels and has been linked to reduced expression of certain NRF2 response genes[44], may be a one of the potential candidates. However, this hypothesis requires further testing in future studies.

Despite the fact that TXNRD1 inhibition shows more pronounced toxicity towards SCLC compared to non-cancerous cells, both in cell culture and mice, we cannot rule out the possibility that dose-limiting toxicity in certain patients may lead to reduced efficacy, even in a susceptible entity like SCLC. However, the observation that SCLCs do not respond to NRF2 induction, while non-cancerous cells become resilient against drug toxicity, enabled us to demonstrate that the therapeutic window for TXNRD1 inhibition can be substantially expanded (Figure 6). Previous studies have shown that organ protection by KEAP1 inhibitors/NRF2 inducers is effective for cisplatin [45] and doxorubicin [46], but a discriminative effect on cancer cells has not been reported before and was, up to this report, counterintuitive.

NRF2 inducers such as CDDO-Me and DMF were primarily developed to treat inflammation associated with chronic diseases (reviewed in [47]) and have shown a toxicity profile that seems compatible with cancer treatments. Furthermore, studies on the immunomodulatory effects of NRF2 pathway activators within the tumor microenvironment (TME) have shown that CDDO-Me attenuates immunosuppression by promoting shifts in macrophage polarization and reducing the numbers of regulatory T cells while simultaneously increasing the number of cytotoxic T cells [48, 49]. These effects lead to more enhanced chemotherapy efficiency in a murine model of lung cancer [50], underscoring the potential of NRF2 activation not only in widening the therapeutic window but also in actively reshaping the immune landscape within the TME.

Having demonstrated that TXNRD1 is a critical vulnerability of SCLC, the question arises whether there are suitable inhibitors available for clinical trials. Auranofin, initially developed for the treatment of rheumatoid arthritis, has been tested repeatedly in clinical trials, but has shown limited success. Based on our data, we attribute this primarily to the lack of appropriate stratification for susceptible, low-ACB tumors. In addition, the molecule itself has physicochemical shortcomings that pose challenges, including poor bioavailability, limited stability, and dose-limiting side effects [51]. A non-metal-containing compound discovered by the Arner lab has shown *in vivo* activity but requires optimization to enhance its stability [52]. DKFZ-608 on the other hand, has shown superior activity and selectivity, compared to auranofin, with favorable bioavailability in mouse models. However, further toxicity and pharmacokinetic studies in non-rodent species are required to provide data for advancing DKFZ-608 into human clinical trials.

In summary, our study demonstrates that low expression of genes involved in ROS-scavenging in SCLC results in the accumulation of lethal ROS levels when a key enzyme of redox homeostasis, TXNRD1, is inhibited. SCLC cells not only experience a rapid exhaustion of their ROS-scavenging capacity but also fail to adapt to oxidative stress by upregulating NRF2-mediated ROS defense.

We conclude that TXNRD1 inhibition is a highly effective strategy for maintenance therapy following cisplatin/etoposide treatment. This is because the resistance mechanisms that emerge in residual cancer cells during first-line therapy do not compromise the effectiveness of TXNRD1 inhibition. Lastly, we propose that combining TXNRD1 inhibitors with NRF2 inducers will maximize the therapeutic potential of TXNRD1 inhibitions in maintenance therapy by protecting healthy tissues and enhancing the selectivity towards cancer cells.

### Supplementary section 1

In HCC33 and H82, in which TXNRD1 inhibition induced high levels of hydrogen peroxide (H_2_O_2_) and hydroxyl radical (HO·) levels, the iron/copper chelator DFO, strongly reduces cell death, presumably by preventing the formation of toxic levels of HO· through Fenton reaction (Supplementary Fig. 4A-C). Interestingly, in those two cell lines, TXNRD1 inhibition caused lipid peroxidation only in HCC33. Ferrostatin-1, which has recently been shown to also act as an iron chelator [53], did not prevent cell death, although it was able to reduce lipid peroxidation. (Supplementary Fig. 4E, F). This suggests that DKFZ-682 induces an iron-dependent form of cell death, distinct from ferroptosis. This type of cell death could not be reduced by inhibitors of necroptosis or AIF- or caspase-dependent apoptosis, indicating catastrophic events leading to cell necrosis (Supplementary Fig. 4D, G). In H1105, where DKFZ-682 causes only a moderate induction of H_2_O_2_ and HO·, toxicity could not be rescued by DFO, pretreatment with olaparib was most efficient to inhibit cell death, indicating the involvement of PARP dependent apoptosis.

### Supplementary section 2

A possible mechanism for the failure to induce ROS buffer genes could be the constitutive nuclear activity of BACH1, a transcriptional repressor of a subset of NRF2 response genes [25, 54]. An earlier report has demonstrated that, in HaCaT cells, CDDO-Me not only increases NRF2-levels but also causes nuclear export of BACH1, leading to a more efficient NRF2 response [55]. Although we couldn’t formally show nuclear export, we detected a reduction of nuclear BACH1 upon CDDO-Me in SCLC cells (Supplementary Fig. 9A, left panel). The combination of the BACH1 inhibitor hemine [56] and CDDO-Me resulted in an increased induction of HMOX1, which has been reported to be positively regulated by NRF2 and repressed by BACH1 [57] (Supplementary Fig. 9A, right panel), confirming that the NRF2/BACH1 regulon is functional in SCLC cells. However, CDDO-Me/hemine pre-treatment did not enable SCLC cells to develop significant resistance against DKFZ-682 (Supplementary Fig. 9B), arguing that continual transcriptional repression by BACH1 is not responsible for the low-ACB status and high drug sensitivity of SCLC cells. We also tested the hypothesis that SCLC achieve low-ACB expression through the downregulation of MAFG, a positive acting binding partner of NRF2 [26]. In Beas-2B, but not in HaCaT cells, knock down of MAFG prevented resistance induction by CDDO-Me, suggesting that ROS buffer capacity in these cells is dependent on genes regulated by NRF2/MAFG heterodimers (Supplementary Fig. 10A). We noted that in SCLC cell lines the MAFG promoter is hyper-methylated in regions where ARE consensus sequences have been predicted (Supplementary Fig. 10B). As MAFG itself is a target of NRF2, it is plausible that its epigenetic repression in SCLC prevents a feed forward loop of ROS response, which had previously been predicted [58]. We circumvented epigenetic silencing of MAFG by tet-inducible overexpression. Our own data and a previous report [59] showed that excessive overexpression of MAF proteins can lead to the repression of NRF2 response genes. We therefor used doxycycline concentrations which allowed a robust but low MAFG induction and found, unexpectedly, that overexpression of MAFG did not enable SCLC cells to develop drug resistance upon NRF2 induction (Supplementary Fig. 10C, D). This suggests that MAFG is not a limiting factor for the defense against drug induced ROS stress in SCLC cells.

---

**Supplementary Figure 1:**
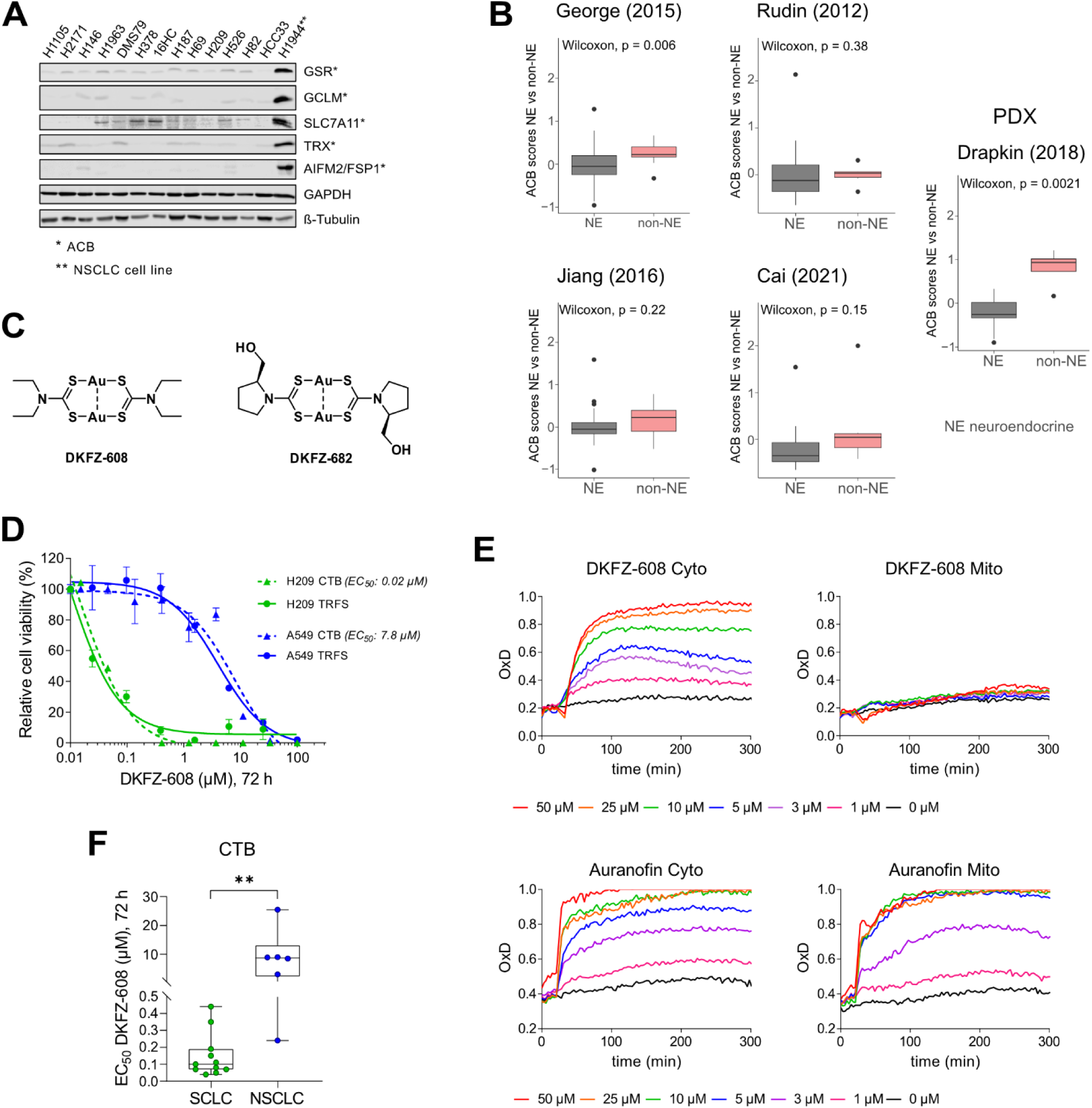
SCLCs express low levels of ACB proteins independent of the neuroendocrine subtype. **(A)** Protein levels of selected ACBs in total cell extracts from 13 SCLC cell lines were analyzed by immunoblotting (n=2). High ACB protein expression in drug resistant H1944 NSCLC cell line was used as a reference. **(B)** The ACB status of SCLC cells derived from PDX models and patient samples showing neuroendocrine (NE) and non-NE characteristics. Expression data were downloaded from the Dryad Digital Repository (https://datadryad.org/stash/share/BkmPdMrwhae1VxhkkSLIG_532FLCqcYiMFUpY1yKmGA) [60]. SCLC samples were categorized into four expression-based subtypes (ASCL1, NEUROD1, POU2F3 and YAP1) based on the highest expression level of key transcription factors. These subtypes were further grouped into “neuroendocrine” (ASCL1 and NEUROD1) and “non-neuroendocrine” (POU2F3 and YAP1) categories to enable comparative analyses between these biologically distinct groups. ACB scores were calculated by taking the average of the scaled expression values for the ACB-genes. Box plots show the median and the interquartile range, and whiskers extend to 1.5x the interquartile range. Significance was determined using the Wilcoxon test. **An impact of DKFZ-608 (TXNRD1 inhibitor) on cell viability and cellular redox status. (C)** Chemical structure of DKFZ-608 and DKFZ-682, previously reported as gold (I)-dithiocarbamate (dtc) complex 5 and complex 37 respectively [17]. **(D, F)** SCLC and NSCLC cell lines were treated with a range of concentrations of DKFZ-608 for 72 h. Cell viability was assessed using CellTiter-Blue (CTB) assay (\*\**p* < 0.01, two-tailed unpaired *t*-test). **(D)** For quantification of intracellular TXNRD1 inhibition, TRFS was added to cell lines, immediately followed by a dilution series of DKFZ-608. Fluorescence induction was measured for 8 h and dose response effects were analyzed at time point 100 min. All data were normalized (untreated control was set to 100 %). Data are presented as mean ± SD of three technical replicates. **(E)** H838 cells expressing either cytoplasmic (Cyto) or mitochondrial (Mito) roGFP2-Orp1 were treated with various concentrations of DKFZ-608 or auranofin. Cells were treated with sulphobuthylester-ß-cyclodextrin as a solvent control for DKFZ-608 and with DMSO as a control for auranofin (black lines). The fluorescence signals from oxidized and non-oxidized cytoplasmic or mitochondrial roGFP2-Orp1 were monitored for 300 min. The oxidation degree (OxD) represents the extent of probe oxidation, with data normalized to 1.0 (fully oxidized) defined by the signal from diamide (2 mM) and 0.0 (fully reduced) defined by the signal from DTT (10 mM). Results are representative of two independent experiments each performed in triplicate.

**Supplementary Figure 2:**
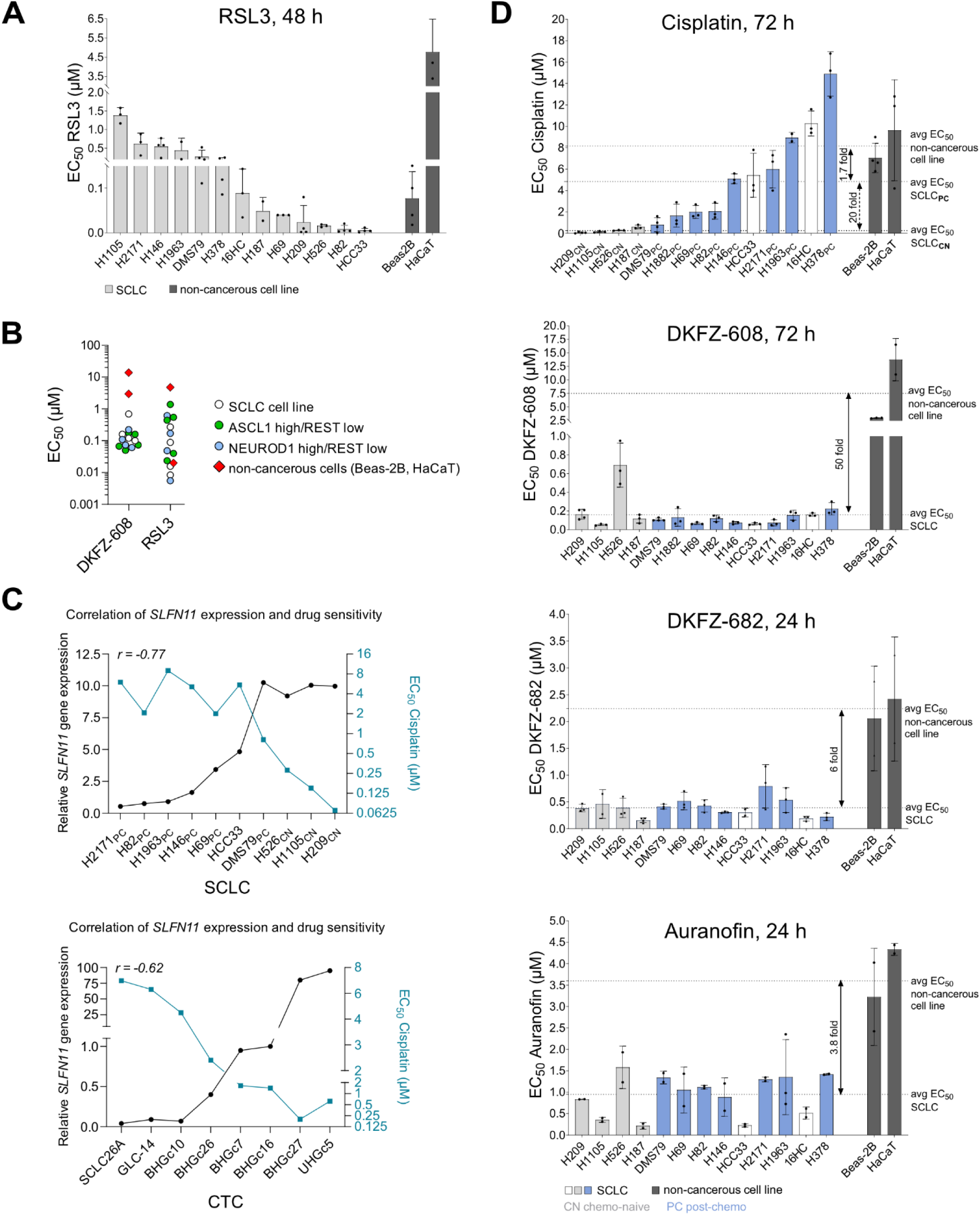
The cytotoxic effect of TXNRD1 inhibition is equal high for all tested SCLC cell lines, independent of their resistance to 1S3R-RSL3 (RSL3) or cisplatin. **(A-D)** SCLC and non-cancerous cell lines were treated with different concentration of drugs for the indicated time point and the cell viability was quantified by the CellTiter-Glo assay. Cells were treated with DMSO as a control for RSL3, cisplatin and auranofin, and with sulphobuthylester-ß-cyclodextrin as a solvent control for DKFZ-608 and DKFZ-682. Bar diagrams show the mean ± SD of EC50 data from independent experiments (n=2-6) each performed in triplicate. **(B)** NE scores were derived from the Gazdar Small Cell Lung Cancer Neuroendocrine Explorer [61]**. (C)** The data for SLFN11 gene expression in SCLC cell lines and circulating tumor cells (CTC) are from DepMap and Gerhard Hamilton respectively (r, Pearson correlation).

**Supplementary Figure 3:**
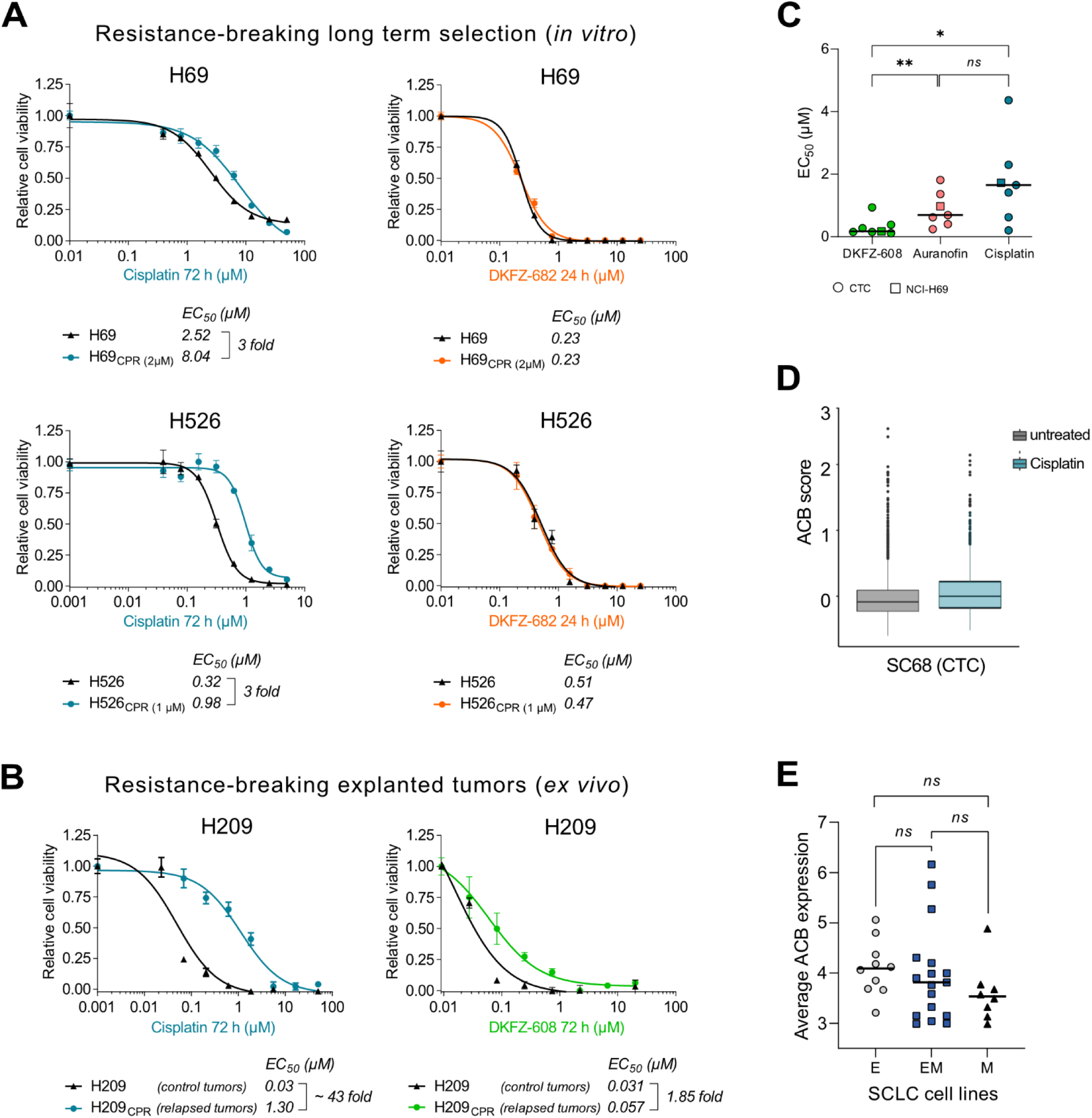
Cisplatin-resistant cells remain sensitive to TXNRD1 inhibition. **(A)** Parental H69 and H526 and cisplatin-resistant (CPR) H69 (resistant to 2 µM of cisplatin) and H526 (resistant to 1 µM of cisplatin) cell lines were exposed to cisplatin for 72 h or DKFZ-682 for 24 h. The cell viability was quantified by the CellTiter-Glo assay. Data represents the mean ± SD of three technical replicates from one of two independent experiments. **(B)** H209 cells, isolated from an etoposide/cisplatin-resistant tumor after 3 cycles *in vivo* therapy, and naïve H209, isolated from control tumors, were treated with various concentrations of cisplatin or DKFZ-608 for 72 h. Surviving cells were quantified by CellTiter-Blue assay. Data are presented as mean ± SD of three technical replicates. **(C)** SCLC H69 cell line and circulating tumor cells (CTC) were seeded in 96 well plates as two-dimensional cultures. EC_50_ values were determined from dose response curves after 96 h treatment with the indicated drugs using a modified MTT assay. Each dot represents a mean of three technical replicates (*ns*, not significant, \**p* < 0.05, \*\**p* < 0.01, ratio paired *t*-test). **Circulating tumor cells (CTC) maintain low ACB expression after cisplatin therapy. (D)** CTCs from naïve and relapsed patients. Data from Stewart et al. 2020 were obtained from the GEO database (accession number GSE138474) and processed by sub-setting and normalizing data from SC68 to focus on epithelial cells. Log counts values were scaled, and the expression levels of ACB genes were averaged to calculate the ACB score. Vehicle-treated and cisplatin-treated cells are compared to assess the impact of treatment. Box plots were used to display the data, with the median and interquartile range shown, and whiskers extending to 1.5 times the interquartile range. Statistical significance was evaluated using the Wilcoxon test. **SCLC cells with distinct profiles of EMT markers demonstrate non-significant differences in ACB expression. (E)** 34 SCLC cell lines were separated according to pre-calculated EMT signatures [62] into epithelial (E), mesenchymal (M) and mixed groups (EM). Based on paired *t*-tests, differences of ACB expression (average values of log2 (TPM+1) converted reads) between groups are non-significant (*ns*).

**Supplementary Figure 4:**
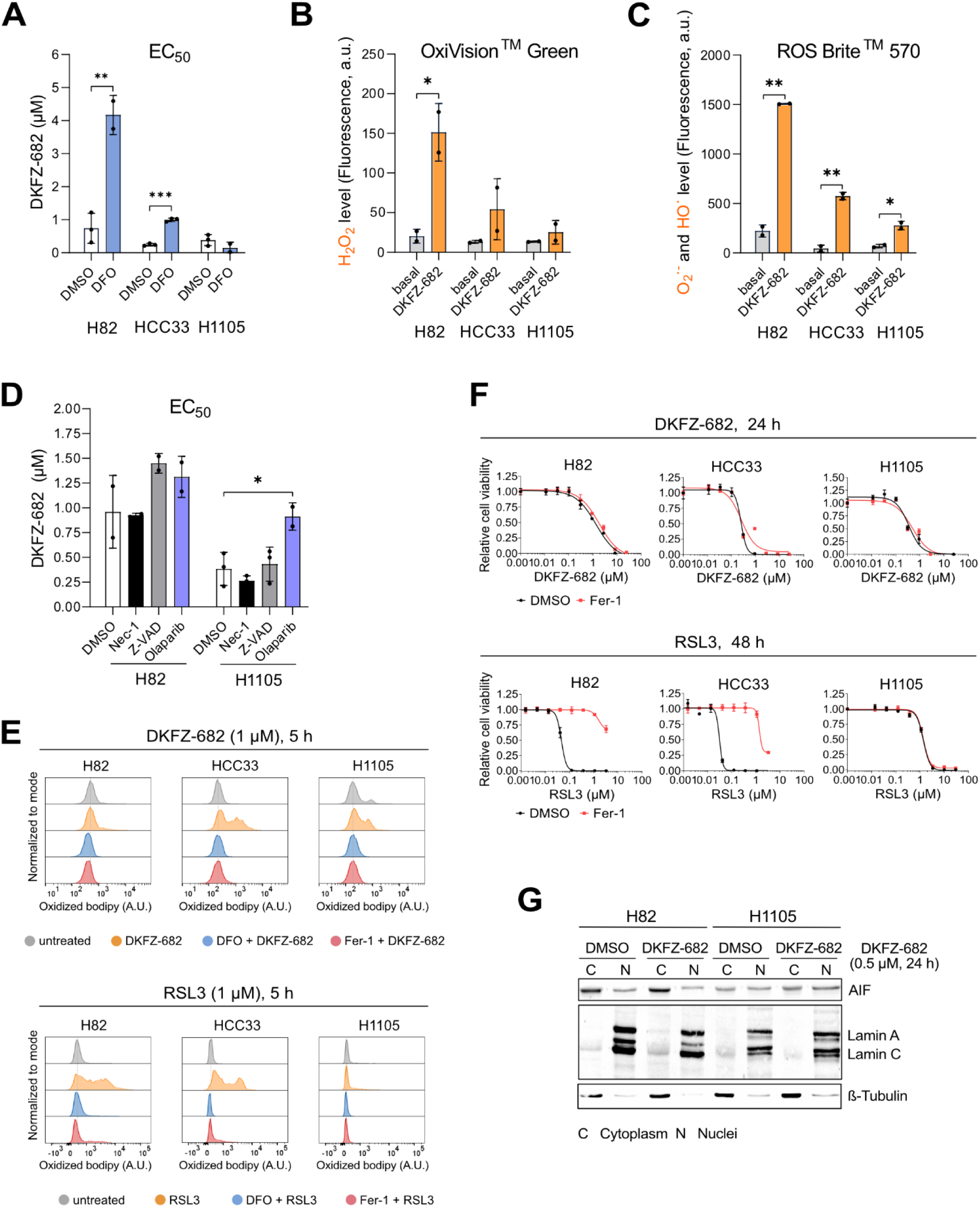
Cell death mechanisms induced by TXNRD1 inhibition in SCLC. **(A)** Cell lines were pre-treated with DMSO or DFO (100 µM) for 1 h. Then the cells were treated with a concentration series of DKFZ-682 for 24 h and cell viability was measured by the CellTiter-Glo assay (EC_50_ mean ± SD, n=2-3; \*\**p* < 0.01, \*\*\**p* < 0.001, two-tailed unpaired *t*-test). **(B, C)** Indicated cell lines were stained with the OxiVision^TM^ Green peroxide sensor or ROS Brite^TM^ 570 fluorescent dye upon treatment with DKFZ-682 (20 µM, 30 min) and analyzed by flow cytometry. Data are presented as mean ± SD of two independent experiment (\**p* < 0.05, \*\**p* < 0.01, two-tailed unpaired *t*-test). **(D)** Cells were pre-treatment with DFO (100 µM) or ferrostatin-1 (Fer-1, 5 µM) for 1 h and treated with DKFZ-682 and RSL3 for 5 h. To analyze lipid peroxidation, cells were stained with bodipy 581/591 C11 and analyzed by flow cytometry. The displayed graphs are representative of 2 independent experiments. **(E)** Cell viability of indicated cell lines pre-treated with Fer-1 (5 µM) for 1 h and treated with DKFZ-682 for 24 h or RSL3 for 48 h was analyzed by CellTiter-Glo assay. The graphs are representative of two independent experiments each performed in triplicate. **(F)** H1105 and H82 cell lines were pre-treated with DMSO, necrostatin-1 (Nec-1, 10 µM), Z-VAD-FMK (20 µM) or olaparib (100 nM). Then the cells were treated with a concentration series of DKFZ-682 for 24 h and the cell viability was measured by the CellTiter-Glo assay. Data represent the mean ± SD of two to four independent experiments, each performed in triplicate (\**p* < 0.05, two-tailed unpaired *t*-test). **(G)** The cells were treated with DMSO (control) or DKFZ-682 for 24 h, and the protein levels of apoptosis inducing factor (AIF) in the cytoplasmic (C) and nuclear (N) fractions were analyzed by Western blotting. Lamin A/C und ß-Tubulin served as controls for the nuclear and cytoplasmic sub-compartments. The results are representative of two independent experiments.

**Supplementary Figure 5:**
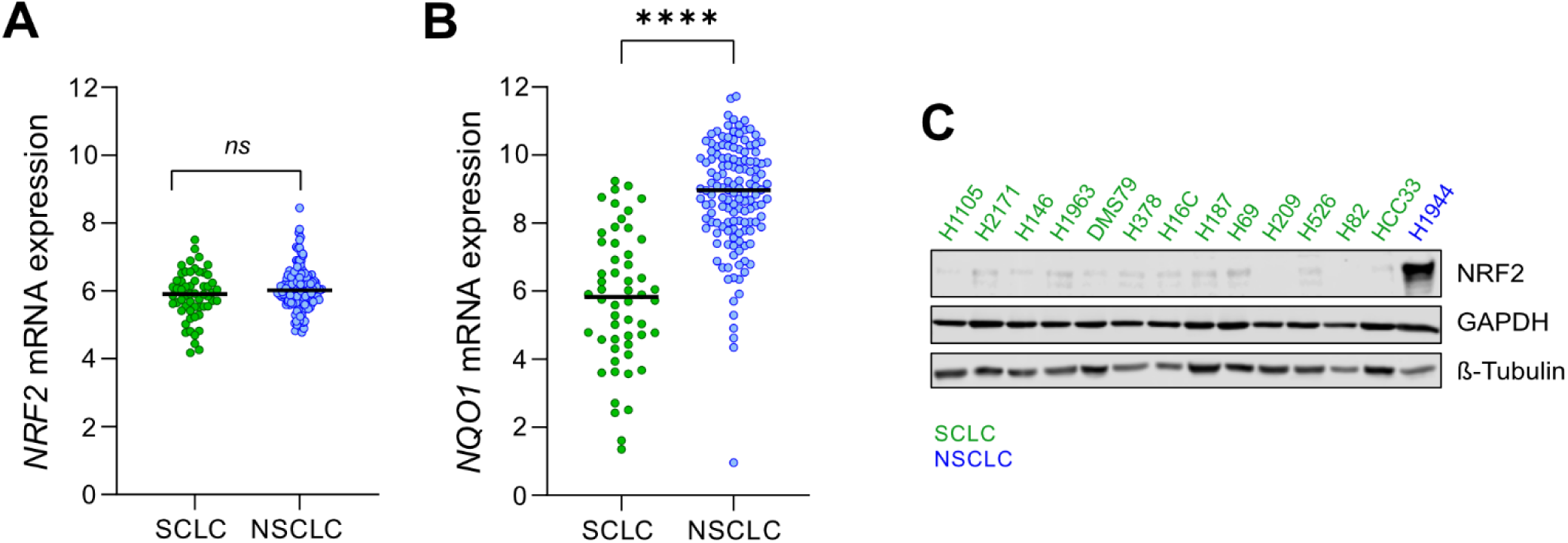
NRF2 transcript levels do not differ between SCLC and NSCLC. **(A)** NRF2 gene expression (data from DepMap) in cells grouped according to their origin (SCLC, NSCLC) does not differ significantly between SCLC and NSCLC (*ns*, not significant, two-tailed unpaired *t*-test). ***NQO1*, a cell type independent proxy for NRF2 activity is expressed at lower levels in SCLC. (B)** *NQO1* gene expression data are from DepMap (**** *p* < 0.0001, two-tailed unpaired *t*-test). **SCLC cell lines express lower levels of NRF2 protein, compared to the KEAP1 mutant cell line H1944. (C)** Representative immune blot (n=2) showing the protein level of NRF2 in total cell extracts from 13 SCLC cell lines. NRF2 expression in drug resistant H1944 NSCLC cell line was used as a reference.

**Supplementary Figure 6:**
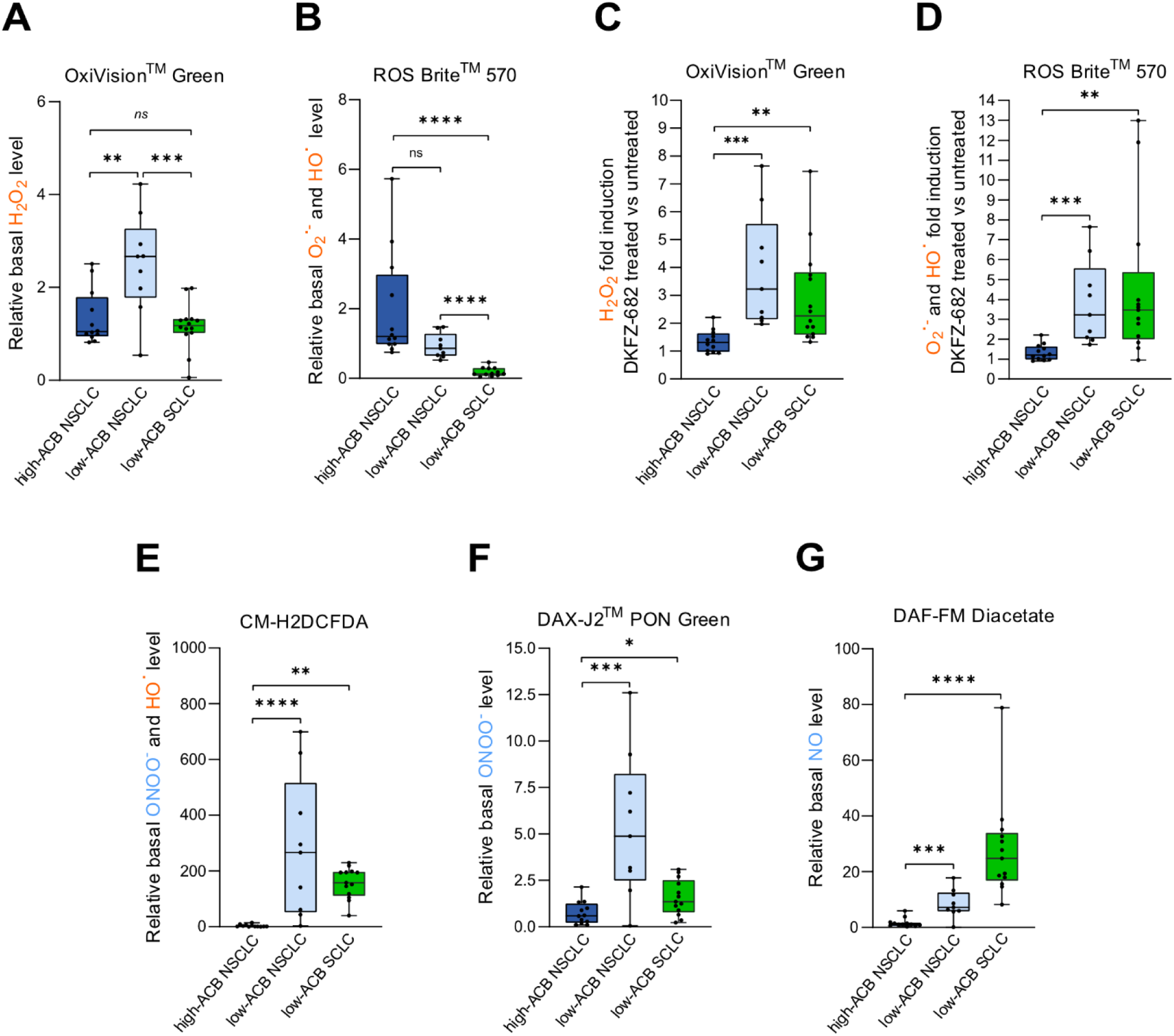
Basal and drug induced ROS and RNS levels in SCLC cell lines. A simplified schematic representation of the main reactive oxygen and nitrogen species (ROS – orange; RNS – blue) and sensors for their detection was described in our previous work [15]. Sensors are nonfluorescent cell-permeant reagents and produce bright fluorescence upon ROS/RNS oxidation. The reagents have good selectivity to H_2_O_2_ (OxiVision^TM^ Green peroxide sensor), O_2_·^−^ (ROS Brite^TM^ 570), HO· (ROS Brite^TM^ 570, CM-H2DCFDA), ONOO^-^ (CM-H2DCFDA, DAX-J2^TM^ PON Green) and NO (DAF-FM Diacetate). The basal ROS **(A, B)** and RNS **(E-G)** levels in high-ACB NSCLC cell lines (n=12), low-ACB NSCLC cell lines (n=9) and low-ACB SCLC cell lines (n=13) were measured by staining with the OxiVision^TM^ Green peroxide sensor **(A)**, ROS Brite^TM^ 570 fluorescent dye [63] **(B)**, CM-H2DCFDA **(E)**, DAX-J2^TM^ PON Green [64] **(F)**, and DAF-FM Diacetate [65] **(G)** and analyzed by flow cytometry. To investigate the changes in ROS levels upon treatment with DKFZ-682 (20 µM for 30 min), cell lines were stained with OxiVision^TM^ Green peroxide sensor **(C)** and ROS Brite^TM^ 570 **(D)**, analyzed by flow cytometry and fold changes were calculated. Each dot (cell line) represents a mean of 2-5 independent experiments (\**p* < 0.05, \*\**p* < 0.01, \*\*\**p* < 0.001, \*\*\*\**p* < 0.0001, two-tailed unpaired *t*-test).

**Supplementary Figure 7:**
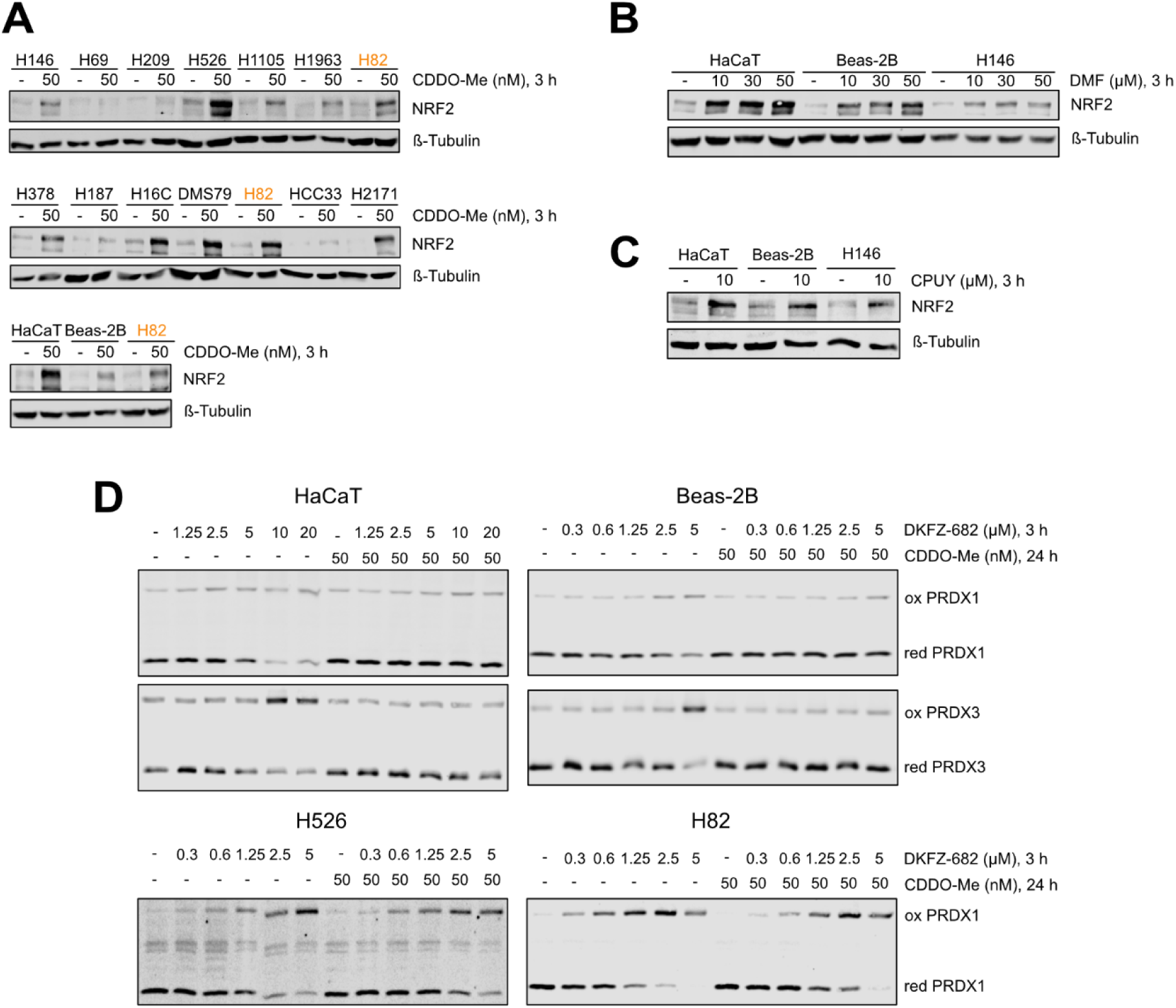
SCLC cell lines remain vulnerable to the cytotoxic effects of drugs, despite their high NRF2 protein level induced by CDDO-Me. **(A-C)** The cells were treated with DMSO (control), CDDO-Me, dimethyl fumarate (DMF) or CPUY192018 (CPUY) for 3 h and the protein level of NRF2 was analyzed by immunoblotting. The blots are representative of two independent experiments. **(D)** Cell lines were pre-treated with CDDO-Me or DMSO for 24 h and then treated with the indicated concentration of DKFZ-682 for 3 h. Oxidized (ox) and reduced (red) levels of PRDX1 and PRDX3 proteins were analyzed by immunoblotting. The blots are representative of at least two independent experiments

**Supplementary Figure 8:**
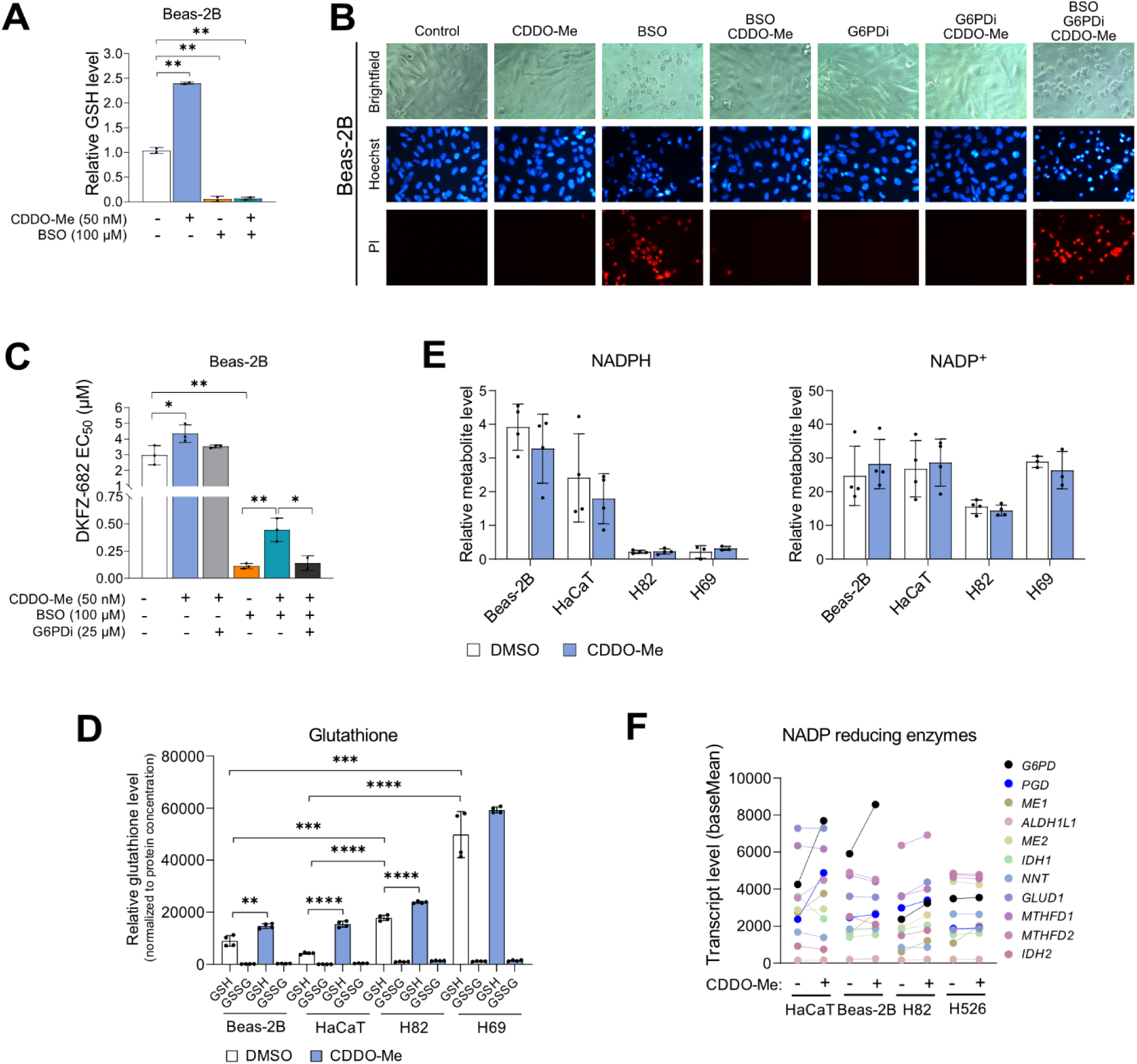
G6PD is involved in CDDO-Me-mediated protection of non-cancerous cells against ROS stress. **(A-B)** The cells were pre-treated with BSO (100 µM) for 24 h, followed by CDDO-Me (50nM) alone or in combination with G6PDi (25 µM) for another 24 h. **(A)** Endogenous levels of reduced glutathione were analysed by reduced glutathione (GSH) assay (Abcam, ab235670; mean ± SD of two biological replicates; \*\**p* < 0.01, two-tailed unpaired *t*-test). **(B)** Cells were co-stained with Hoechst 33342 to visualize all cells and with propidium iodide (PI) to identify dead cells. The phase contrast and fluorescence microscopy images are representative of two independent experiments. **(C)** The cells were pre-treated with BSO for 24 h, followed by CDDO-Me alone or in combination with G6PDi for another 24 h. Then the cells were treated with a concentration series of DKFZ-682 for 24 h. Cell viability was measured by the CellTiter-Glo assay. Data represent the mean ± SD of 3 independent experiments, each performed in triplicate (\**p* < 0.05, \*\**p* < 0.01, two-tailed unpaired *t*-test). **(D)** Cell lines were treated with DMSO (control) or CDDO-Me (50 nM) for 24 h. Endogenous levels of glutathione were analyzed in cell lysates using GSH/GSSG-Glo^TM^ assay (Promega, V6611). The signal intensities were normalized to protein concentration of each sample. The results are mean ± SD of two independent experiments each performed in duplicates (\*\**p* < 0.01, \*\*\**p* < 0.001, \*\*\*\**p* < 0.0001, two-tailed unpaired *t*-test). **(E)** The endogenous levels of NADPH and NADP^+^ were analysed using metabolomic analysis. The signal intensities were normalized to protein concentration of each sample (n=3-4). **(F)** The transcript levels of enzymes involved in NADPH regeneration upon CDDO-Me treatment as determined by the expression profiling analysis (n=3).

**Supplementary Figure 9:**
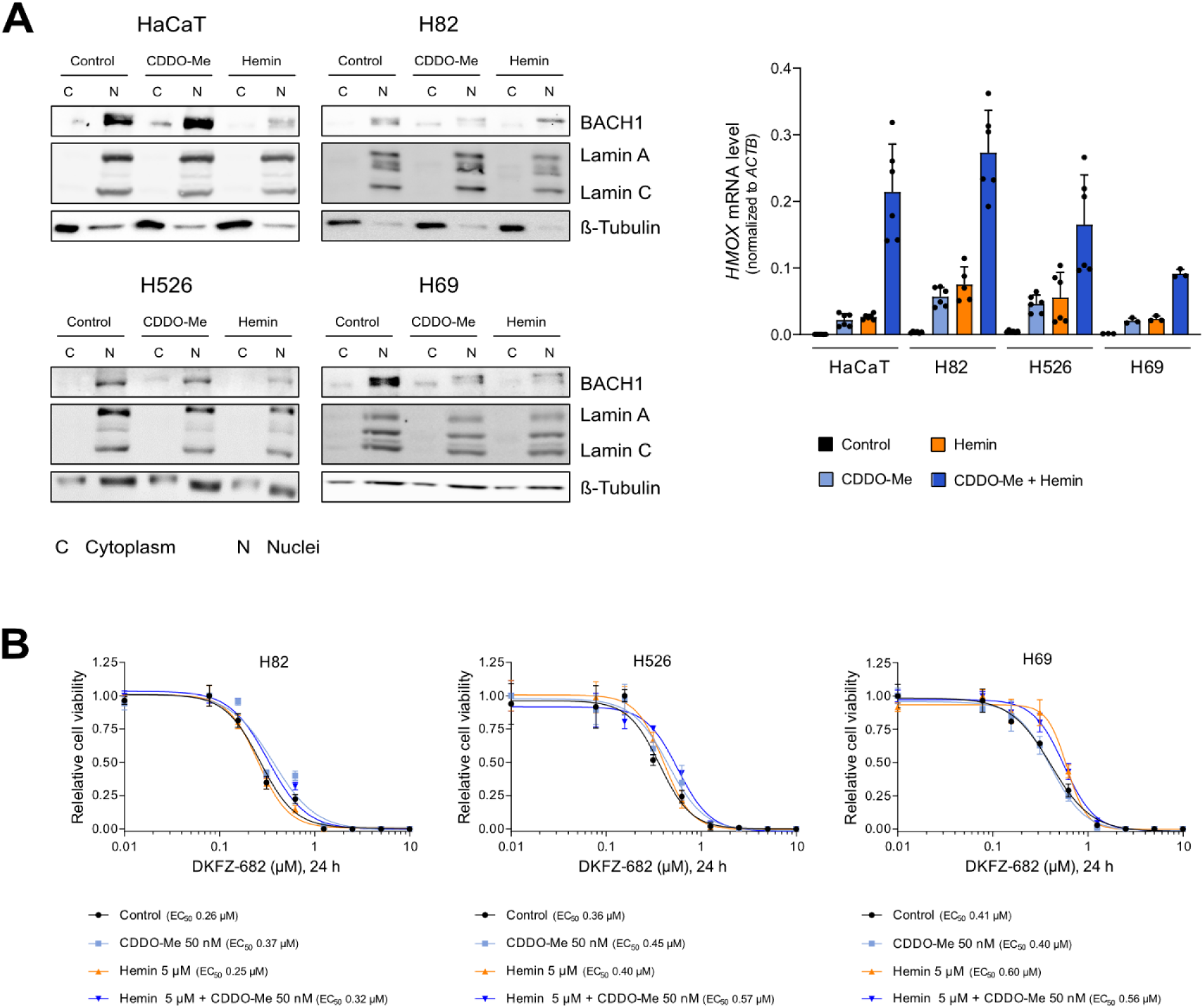
Inhibition of BACH1 does not affect sensitivity of SCLC to TXNRD1-targeting drug. **(A)** The indicated cell lines were treated with DMSO (control), CDDO-Me (50 nM) or hemin (10 µM) for 6 h. The protein levels of BACH1 (left panel) were analyzed in the cytoplasmic (C) and nuclear (N) fraction by immunoblotting. Lamin A/C and β-Tubulin served as controls for the nuclear and cytoplasmic sub-compartments, respectively. The blots are representative of two independent experiments. The level of *HMOX* (right panel), an indicator of BACH1 inhibition, was analysed by qPCR. The results (mean ± SD) are representative of two independent experiments, each performed in triplicates. **(B)** The cells were pre-treated with DMSO (control), CDDO-Me (50 nM) with/or hemin (5 µM) for 24 h, and then treated with dilution series of DKFZ-682 for 24 h. The cell viability was quantified by the CellTiter-Glo assay. The results are representative of two independent experiments, each performed in triplicates.

**Supplementary Figure 10:**
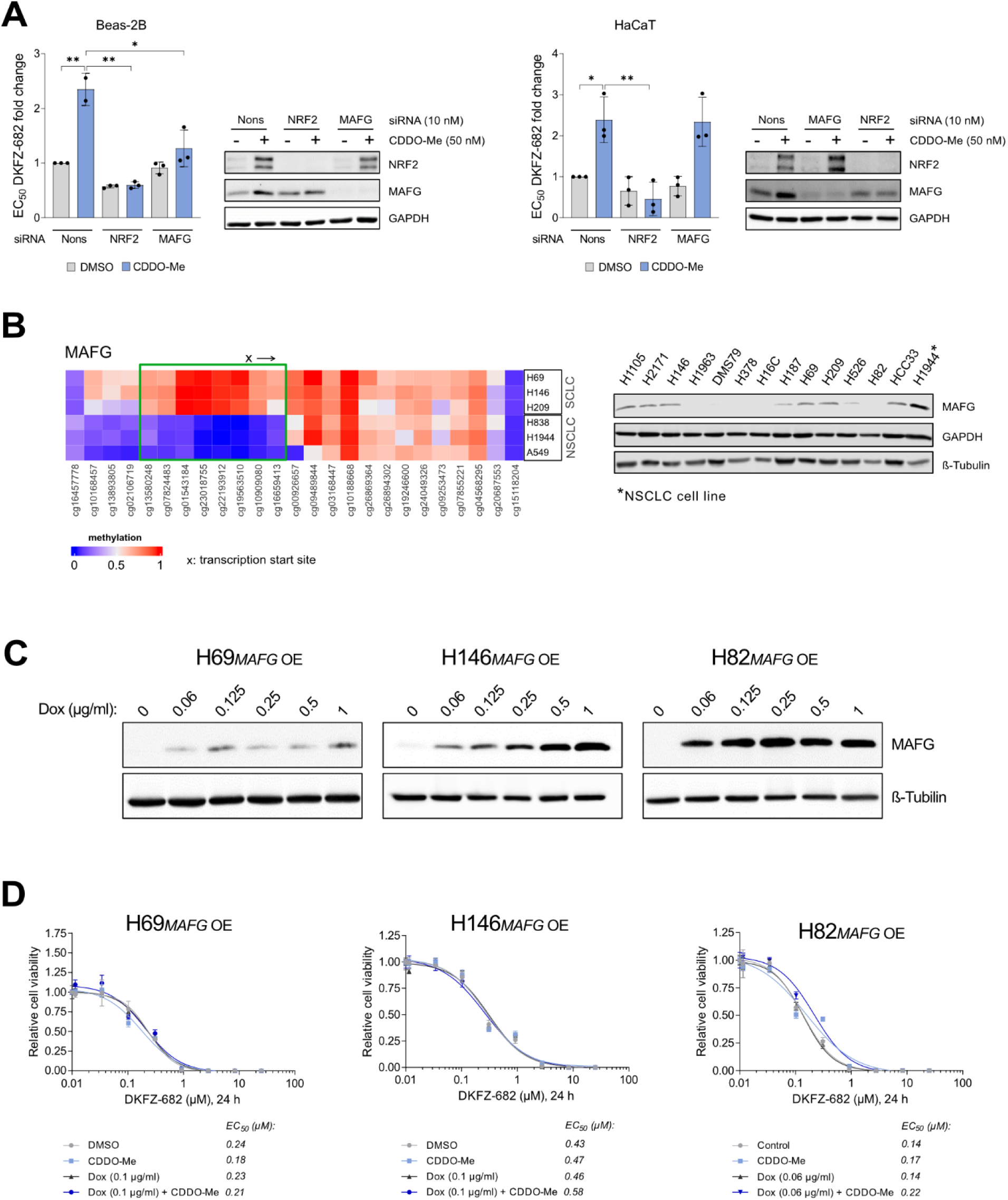
MAFG is essential for CDDO-Me inducible protection of Beas-2B against of cytotoxic effect of TXNRD-1 inhibition. **(A)** Cells were either treated with nonsense (Nons) siRNA or siRNA against *NRF2* or *MAFG*. The next day, cells were trypsinized and 5,000 cells were seeded into the wells of a 96-well plate. After 24 h, cells were treated with DMSO (control) or CDDO-Me (50 nM). After additional 24 h, cells were treated with a range of concentrations of DKFZ-682 for 24 h. Cell viability was assessed using CellTiter-Glo assay. The graph is representative of independent experiments (n=2-3, mean ± SD; \**p* < 0.05, \*\**p* < 0.01 two-tailed unpaired *t*-test) each performed in triplicates. EC_50_ fold change data are normalized to Nons. NRF2 and MAFG protein expression was analysed by immunoblotting. Representative western blots are shown. Increased promoter methylation of *MAFG* gene in SCLC. (B) The heatmap, showing the methylation status of genomic DNA (left panel; 0 – no methylation, 1 – high methylation) was analysed using the Infinium MethylationEPIC BeadChips (Illumina). The protein level of MAFG (right panel) in total cell extracts from 13 SCLC cell lines was analyzed by immunoblotting (n=2). High MAFG expression in drug resistant H1944 NSCLC cell line was used as a reference. **Combined NRF2 induction by CDDO-Me and overexpression of MAFG in SCLC cells does not lead to a desensitization of cells to TXNRD1 inhibition. (C)** Cells with a tet-inducible MAFG expression construct were treated with doxycycline (Dox) for 24 h to increase MAFG expression. The protein level of MAFG was analyzed by immunoblotting. **(D)** Cells were pre-treated with Dox and/or CDDO-Me (50 nM) for 24 h to increase MAFG and NRF2 expression respectively. Then the cells were treated with a concentration series of DKFZ-682 for 24 h and cell viability was measured by the CellTiter-Glo assay. The graph is representative of two independent experiments each performed in triplicate.

**Supplementary Figure 11:**
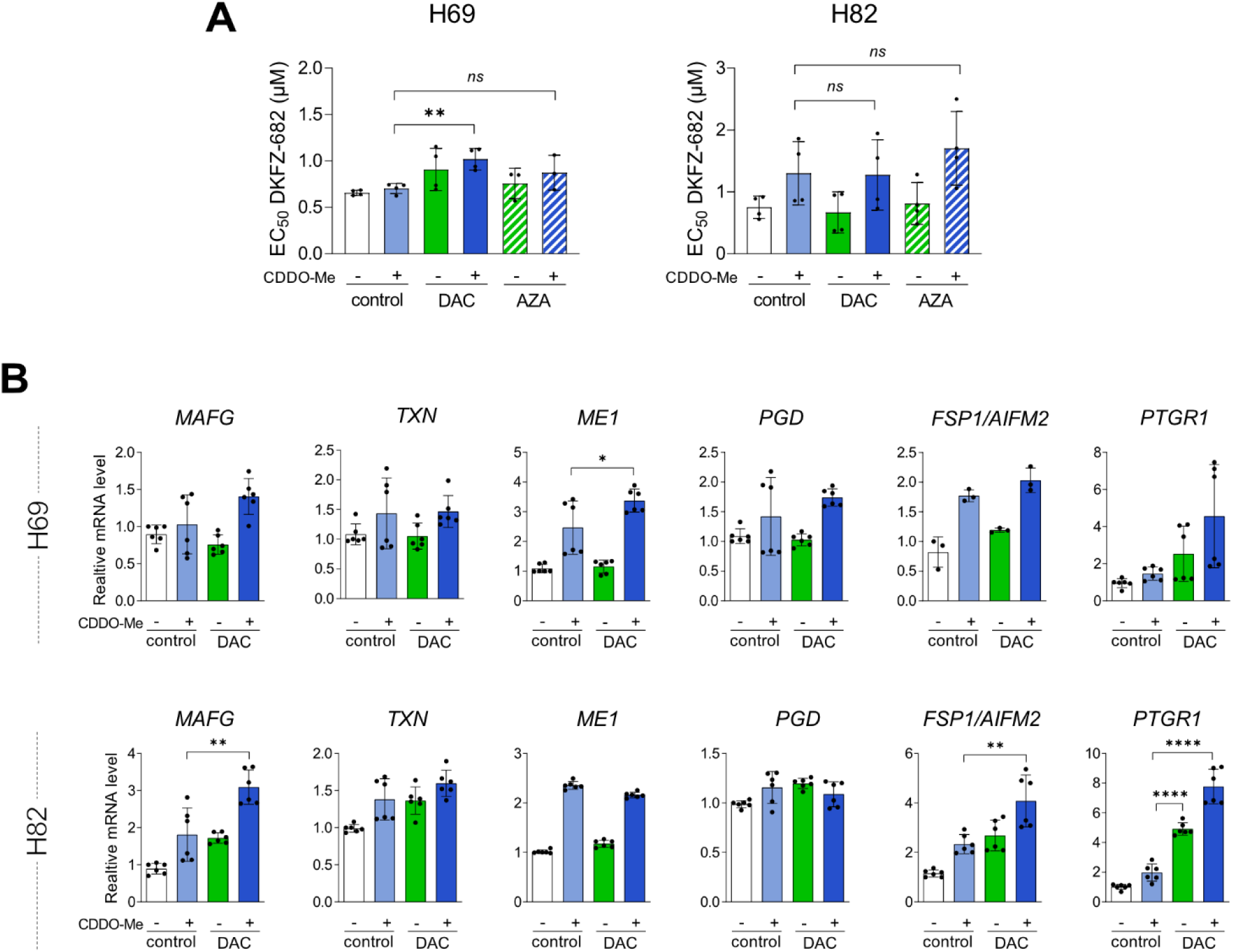
Combined Effect of pre-treatment with DAC and CDDO-Me on cell sensitivity to TXNRD inhibition and on ACB gene expression. **(A, B)** H69 and H82 cell lines were treated with decitabine (DAC, 0.5 µM) or azacitidine (AZA, 2.5 µM) three times over one week (on days one, three, and five). Afterward, the medium was replaced with DAC/AZA-free medium. **(A)** Cells were seeded in a 96-well plate and pre-treated with or without CDDO-Me (50 nM). After 24 hours, the cells were exposed to a concentration series of DKFZ-682 for 24 hours, and cell viability was measured using the CellTiter-Glo assay. Data represent the mean ± SD of independent experiments (n=3-4), each performed in triplicate (\*\**p* < 0.01, two-tailed unpaired *t*-test). **(B)** Cells were treated with CDDO-Me (50 nM) for 6 h and levels of MAFG and ACB transcripts were determined by qPCR (Control without CDDO-Me was set to 1; relative data represent mean ± SD of two independent experiments each performed in triplicate; *ns*, not significant, \**p* < 0.05, \*\**p* < 0.01, \*\*\*\**p* < 0.0001, two-tailed unpaired *t*-test).

**Supplementary Figure 12:**
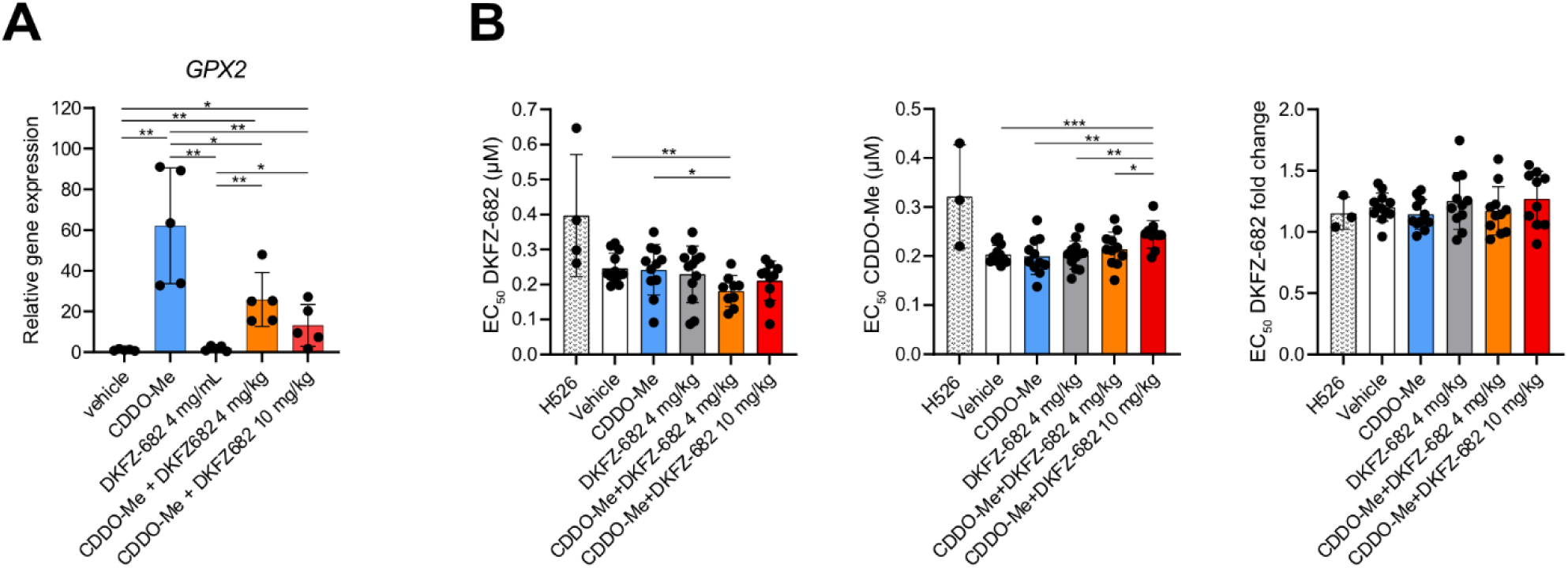
Impact of CDDO-Me pre-treatment on GPX2 expression and DKFZ-682 sensitivity in tumors. **(A)** *GPX2* transcript levels were determined by qPCR in tumors after the end of the experiment in 5 animals from each treatment group (one-way ANOVA *p* < 0.0001; \**p* < 0.05, \*\**p* < 0.01 two-tailed unpaired *t*-test). **(B)** The sensitivity to DKFZ-682 and CDDO-Me was tested in explanted tumors. Tumor cells were treated with a concentration series of DKFZ-682 or CDDO-Me for 24 h and cell viability was quantified using CellTiter-Glo assay. Each dot represents a tumor from one mouse measured in a triplicate. Next, tumor cells were pre-treated with CDDO-Me (50 nM) for 24 h and then the concentration series of DKFZ-682 was applied. The fold change of EC_50_ for DKFZ-682 upon CDDO-Me compared to vehicle pre-treated cells was calculated (one-way ANOVA *p* < 0.0001; \**p* < 0.05, \*\**p* < 0.01, \*\*\**p* < 0.001 two-tailed unpaired *t*-test).

## Materials and Methods

### Synthesis of DKFZ-608

[Au (diethyldithiocarbamate)]_2_ (DKFZ-608): To a solution of aurothiomalate (2.0 g, 5.13 mmol, 1.0 eq.) in H_2_O (100 mL) was added a solution of tetraethyldithiuram disulfide (1.52 g, 5.13 mmol, 1.0 eq.) in EtOH (100 mL). An orange precipitate formed. The suspension was stirred at room temperature for 3 h, the obtained precipitate was filtered through a sintered glass frit (pore size 4), thoroughly washed with water, and dried under high vacuum overnight. The crude product was recrystallized from hot DMF to afford 1.21 g (68 % yield) of DKFZ-608 as orange needles. Elemental analysis [M]: calculated C, 17.39; H, 2.92; N, 4.06; found C, 17.45; H, 3.02; N, 4.22. Analysis of the obtained crystals by x-ray crystallography revealed a molecular unit as previously reported [66]. Due to the low solubility of this compound, a solution of DKFZ-608 (5 mM) was prepared in 150 mM ß-Cyclodextrin sulfobutyl ether/PBS [17] by heating up in a water bath at 100 °C and repeated vortexing every 10-15 min until fully dissolved. The stock solution was stable for more than 2 months at 4 °C and continuously checked for activity in biochemical and cellular assays. In order to use equivalent amounts of gold atoms when comparing DKFZ-608 to auranofin, we calculated concentrations according to gold content.

### Cell culture and siRNA transfection

Human SCLC suspension cell lines (ATCC: NCI-H69, NCI-H82, NCI-H526, NCI-H209, NCI-H1105, NCI-H187, DMS79, NCI-H146, NCI-H2171, NCI-H1963, NCI-H378; DSMZ: HCC33; [67]: SCLC-16HC) and the NSCLC cell line NCI-H1944 (ATCC) were cultivated in RPMI-1640 (Gibco). Other NSCLC cell lines A549 and NCI-H838 (both ATCC) were grown in low-glucose and high-glucose DMEM (Gibco), respectively. The human non-cancerous Beas-2B (ATCC) and HaCaT (AddexBio) were grown in DMEM/F-12 (Gibco) and in high-glucose DMEM (Sigma-Aldrich) medium, respectively. Prior to seeding for experiments, SCLC cells were harvested by centrifugation (4 min at 130 *g* at room temperature). H838, A549, H1944, Beas-2B and HaCaT cells were split by incubation with trypsin-EDTA (0.25 %) for 5 min at 37 °C. All the media contained 10 % fetal bovine serum (FBS-12A, Capricorn Scientific), 100 U/mL penicillin and 100 µg/mL streptomycin (Gibco). MAFG overexpressing SCLC cells were cultured in medium with puromycin (0.5 µg/mL) to make sure that only cells that carry insertion survive. The cells were cultivated at 37 °C in a 5 % CO_2_ atmosphere. Cultures derived from circulating cancer cells were handled as described in [68].

Cell lines were tested negative for *mycoplasma* contamination (Eurofins Genomics). Cell lines were authenticated using Multiplex Cell Authentication by Multiplexion (Heidelberg) as described in [69]. Explanted tumors were stored in a cryopreservation medium (10 % DMSO, 36 % FBS, 54 % RPMI-1640) in liquid nitrogen. After thawing, cells were filtered through EASYstrainer 40 µm (Greiner), washed with PBS and cultivated in RPMI-1640 (Gibco) supplemented with 10 % fetal bovine serum (FBS-12A, Capricorn Scientific), 200 U/mL penicillin and 200 µg/mL streptomycin (Gibco).

Small interfering RNA (siRNA) to knock down target genes were ordered from siTOOLS (Planegg, Germany). Transient transfection was performed using Lipofectamine® RNAiMAX Transfection Reagent according to the manufacturer’s instructions (Thermo Fisher Scientific, Germany).

### Chemical compounds

Chemical compounds used in this study are listed in **Supplementary Table S1**.

### Generation of MAFG overexpressing cells

Sequences of primers used for cloning are provided in **Supplementary Table S2**. The ORF of human *MAFG* was PCR amplified from plasmid carrying this transgene with primers containing SalI and AvrII restriction sites using Phusion High-Fidelity Polymerase (Thermo Fischer Scientific) and inserted into the lentiviral pLIX_403 vector. The sequence of construct was confirmed by sequencing (Microsynth).

Lentivirus was produced through transfection of lentiviral backbones containing transgene (pLIX-MAFG) along with third-generation packaging plasmids into HEK293T cells according to the Trono laboratory protocol [70]. Fresh lentivirus from transfected HEK293T culture supernatant was used for viral transduction, specifically 300 µL of fresh lentivirus was added to 1 million of suspension cells (H82 and H69). All experimental procedures for lentivirus production and transduction were performed in a biosafety level 2 laboratory. 24 h after transduction, lentivirus was removed from the media by centrifugation (300 *g*, 3 min) and cells were kept with 3 mL of fresh medium in 6-well plates for 48 h. Puromycin (0.5 µg/ mL) was added to cells for antibiotic selection for one week. Puromycin resistant cells were kept without puromycin to recover and used for further experiments. The cells armed with a tet-inducible MAFG expression construct were treated with doxycycline (0.06 −1 µg/mL for 24 h) to increase MAFG expression and the level of protein was validated by Western blotting.

### Generation of cisplatin-resistant cells

Cisplatin-resistant (CPR) SCLC cell lines were established from each original parental cell line through continuous exposure to cisplatin over a period of approximately 6 months. Each cell line (2×10^6^ cells in 6 mL culture medium per flask) was treated with freshly prepared cisplatin for 72 h. The initial cisplatin concentration was individually optimized for each cell line (e.g., 0.25 or 0.5 µM for H69 and 0.1 µM for H526 cell lines). Following treatment, the medium was replaced, and the cells were allowed to recover in fresh, drug-free medium for an additional 96 h. The treatment was repeated sequentially until the cells demonstrated adaptation to the currently treated drug concentration. Adaptation was assessed by observing changes in the color of the culture medium (from pink to yellow, indicating culture medium consumption) and an increase in cell aggregate size. Upon confirmation of adaptation, the drug concentration was incrementally increased. For the experiments, cells cultured for 2 weeks in cisplatin-free medium were used. Cell resistance to the final cisplatin treatment concentration (2 µM for H69 and 1 µM for H526) was confirmed using CellTiter-Glo assay.

### Cell viability assays and determination of EC_50_

Cells were seeded in white 96-well plates at a density of 5,000 cells/well in 100 µL of medium. After 24 h, test compounds were added (all concentrations in triplicate) without medium change. After an incubation for the indicated time period, viable cells were quantified using either CellTiter-Glo or CellTiter-Blue cell viability assay (Promega) according to the manufacturer’s instructions. EC_50_ values were calculated from dose response curves by GraphPad Prism.

Circulating SCLC cells were seeded at a density of 10,000 cells in 100 µL of medium in transparent 96-well plates. Test compounds were added in triplicate and the plates were incubated for four days under tissue culture conditions and viable cells detected using a modified MTT assay (EZ4U, Biomedica). EC_50_ values were determined using Origin 9.1 software (OriginLab).

### Monitoring of TXNRD1 activity in cells with TRFS-Green

The probe TRFS-Green was synthetized according to Zhang et al. [71]. Cells (7,500 cells/well) were seeded in V-bottom 96-well plates in FluoroBrite DMEM medium (Gibco) and incubated at 37 °C in 5 % CO_2_ atmosphere. After four hours, DKFZ-608 was added at indicated concentrations in triplicates, and TRSF-Green (10 µM) was immediately added. Following mixing with a multi-channel pipette, plates were centrifuged at 200 *g* for 5 min. Fluorescence (450/520 nm) was monitored using a CLARIOstar plate reader at 2 min intervals over 8 h at 37 °C. Dose-response curves were generated by calculating the fluorescence increase during the linear phase (e.g. 100 min) for each well. Mean values +/- SD were plotted.

### Gel electrophoresis and Western blotting

For total protein extraction, cells were harvested using Pierce^TM^ RIPA buffer (Thermo Scientific, 89900) supplemented with a protease inhibitor cocktail (Serva), 100 U/ml benzonase (Merck) and, if necessary, PhosSTOP phosphatase inhibitor cocktail (Roche). Protein concentrations were measured by Nanodrop (Thermo Fisher).

Preparation of cytoplasmic and nuclear extracts was performed as described previously [15]. Tissue homogenates were prepared from tissue pieces stored at −80 °. Samples were quickly weighed and ice-cold PBS supplemented with the protease inhibitor cocktail was added to each sample to make 10 % homogenate while using TissueLyser II (Qiagen) with one stainless steel bead (5 mm) per sample (4x 30 s at 25 Hz). Protein concentration was determined by DC Protein Assay (Bio-Rad) using BSA as a standard.

Cell lysates and tissue homogenates were denatured by incubation with Laemmli sample buffer (5.1 % glycerol, 0.51 % SDS, 0.051% bromophenol blue, 30 mM DTT, 10.6 mM Tris-HCl, pH 6.9) for 5 min at 95 °C. Proteins were separated with Tris-glycine SDS-PAGE, transferred onto a nitrocellulose membrane, and detected with primary antibodies according to manufacturers’ instructions, followed by incubation with secondary antibodies conjugated to fluorophores. Fluorescence was recorded with either the Odyssey Sa Imager (Li-Cor) or the Chemidoc Imager (Bio-Rad) and quantified using the Image Studio Lite (Li-Cor) or the Image Lab software (Bio-Rad), respectively. The antibodies used in this study are listed in **Supplementary Table S3**.

To detect oxidized PRDX1 and PRDX3, cell lysates were prepared as described previously [15].

### DNA methylation analysis of cell lines

Genomic DNA extracted from 3 SCLC lines (H209, H146 and H69) and 3 NSCLC lines (H838, H1944 and A549) was subjected to methylation analysis using the Infinium MethylationEPIC BeadChips (Illumina) allowing the simultaneous quantitative measurement of the methylation status at 865,918 CpG sites. Approximately two million cells were harvested either by centrifugation at 1,000 *g* for 5 min (suspension cultures) or by scraping from the dishes, followed by centrifugation at 1,000 *g* for 5 min (adherent cells).

### DNA methylation analysis of patient samples

FlexiGene DNA Kit (Qiagen) was used for isolation of DNA according to the manufacturer’s instructions. No technical replicates were performed. DNA concentrations were determined using PicoGreen staining (Molecular Probes). The quality of genomic DNA samples was checked by agarose-gel analysis, and samples with an average fragment size >3kb were selected for methylation analysis. The laboratory work was done in the Genomics and Proteomics Core Facility at the German Cancer Research Center, Heidelberg, Germany (DKFZ). 500 ng of genomic DNA from each sample was bisulfite converted using the EZ-96 DNA Methylation Kit (Zymo Research) according to the manufacturer’s recommendations. The DNA was applied to Infinium MethylationEPIC BeadChip and hybridization was performed for 16-24 h at 48 °C. Allele-specific primer annealing was followed by single-base extension using DNP- and Biotin-labeled ddNTPs. After extension, the array was fluorescently stained, scanned, and the intensities at each CpG were measured. Microarray scanning was done using an iScan array scanner (Illumina). DNA methylation values, described as beta values, were recorded for each locus in each sample. Data analysis was performed with the Illumina’s GenomeStudio 2011.1 (Modul M Version 1.9.0). The complete raw Illumina data was quantile normalized.

### Methylation heatmaps based on CpG clusters

We plotted methylation values for CpG clusters (file CCLE_RRBS_TSS1kb_20181022.txt.gz from DepMap) within promoters of ACB gene set (and MAFG) using ComplexHeatmap R package for cell lines from DepMap project (CCLE) that have both promoter methylation and expression data (Figure 4A). We calculated correlation between vector of cell lines’s methylation values and vector of cell lines’s average expression of genes from ACB set per each CpG cluster, and plotted these correlation values as heatmap’s column annotation. As heatmap’s row annotation we plotted average expression of genes from ACB gene set per cell line and sorted heatmap’s rows according to this value (from highest avg. ACB expression to the lowest). We also provided an additional layer of row annotation – cell line’s EC_50_ values, and additional layer of column annotation – correlation between vector of cell lines’ methylation values and vector of cell lines’ EC_50_ per each CpG cluster.

### Immunohistochemistry (IHC)

The IHC staining in human samples was supported by the tissue bank of the National Center for Tumor Diseases (NCT, Heidelberg) in accordance with the ethical regulations of the NCT tissue bank established by the local ethics committee. IHC was performed by applying a rabbit monoclonal antibody against the TXN1 (1:2000; Abcam, ab133524) to tissue microarrays (TMA, two cores per case) of formalin fixed and paraffin embedded (FFPE) NSCLC and SCLC specimens. The staining procedure was carried out using an autostainer (Bench-Mark ULTRA, Ventana Medical Systems) according to the manufacturer’s instructions.

Staining intensities were evaluated digitally using the QuPath software (v. 0.1.2) [72] and applying random tree algorithms for classification. After verification by a pathologist, the classifiers were used to differentiate between non-tumor cells (e.g. fibroblast, lymphocyte) and tumor cells and to count the latter subdivided by three staining intensities (weak, moderate, strong) in each TMA core. For a weighted analysis, we calculated H-scores, yielding from 0 to 300 as the sum of the percentage of weakly, moderately and strongly stained tumor cells multiplied by 1, 2 and 3, respectively. We averaged the H-scores over the two TMA cores of each case.

### RNA isolation and RT-qPCR

Total RNA from cells was isolated using NucleoSpin RNA Kit according to the manufacturer’s protocol (Macherey-Nagel). To isolate RNA from mouse tissues stored in RNAlater (Sigma-Aldrich), samples were first homogenized in the TRIzol reagent (Invitrogen) using TissueLyser II (Qiagen) with one stainless steel bead (5 mm) per sample (2x 2 min at 25 Hz) and then the manufacturer’s instructions were followed. Purified RNA was measured by Nanodrop (Thermo Fisher) and reverse transcription was performed using Revert Aid First Strand cDNA Synthesis Kit (Thermo Scientific). qPCR was performed on the Roche Lightcycler 480 system using Blue S’Green qPCR kit (Biozym). Primer sequences used for quantitative real-time PCR analyses are listed in **Supplementary Table S4.** *GAPDH* or *ACTB* was used for normalization of gene expression.

### ACB expression profile (Figure 1C)

Expression data for normal tissues, lung adenocarcinoma (LUAD) and lung squamous cell carcinoma (LUSC) were downloaded from The Cancer Genome Atlas (TCGA) and converted to Log2(TPM+1) values. SCLC expression values were obtained from [11] and also converted to Log2(TPM+1) values. All expression values were then z-scored and normalized to normal lung tissue. To determine the ACB scores, the expression of ACB genes was averaged. The relative ACB expression levels in normal tissues, lung squamous cell carcinoma (LUSC), and lung adenocarcinoma (LUAD) were compared to small cell lung cancer (SCLC) using the Wilcoxon test.

Expression profiles for pulmonary neuroendocrine cells (PNECs) were obtained from the Human Lung Cell Atlas through Synapse (https://www.synapse.org/#!Synapse:syn21041850) [73]. Smart-seq2 single-cell profiles from PNECs were aggregated by patient and processed as described above.

### Expression profiling (cell lines)

Total RNA isolated from Beas-2B, HaCaT, H82 and H526 cells using NucleoSpin RNA Kit according to the manufacturer’s protocol (Macherey-Nagel) was submitted to transcriptome profiling on Human Clariom S Assay chips (Applied Biosystems) that allow for quantification of >20,000 well-annotated genes. All the necessary quality controls and the analysis itself were performed by the Microarray Core Facility of the German Cancer Research Center.

### RNAseq (mouse tissues and tumors)

Total RNA extracted from mouse tissues and tumors was submitted for RNAseq analysis to the NGS Core Facility of the German Cancer Research Center. After RNA passed the quality control, sequencing libraries were prepared using TruSeq Stranded mRNA Library Prep Kit (Illumina) and IDT Unique Dual Indexes for Illumina (Integrated DNA Technologies) and sequenced using NovaSeq 6000 sequencing system (paired-end mode, read length 100 bp, S4 flow cell; Illumina), yielding on average 52 million reads per sample (32–76 million). For the analysis of the known reference transcriptome assembly, RNA sequencing data were processed automatically via the One Touch Pipeline (OTP) [74].

### Animal experiments

All studies involving mice were conducted in compliance with German Cancer Research Center guidelines and approved by the governmental review board of the state of Baden-Württemberg, Karlsruhe District Council, under authorization no. G-191/16, G-259/18, and G-176/19, according to German legal regulations. Mice were housed in individually ventilated cages under temperature and humidity control. Cages contained an enriched environment with bedding material. Animal health was monitored daily and mice were euthanized as soon as they reached the termination criteria defined in the procedure. Sample size was calculated with the help of a biostatistician. Assumptions for the power analysis were as follows: Alpha error, 5%; Beta error, 20%. Mice were randomized into treatment groups before treatment. In case animals had to be euthanized before the predefined end point (due to weight loss or other termination criteria), they were excluded from any downstream analyses.

The solution of DKFZ-608 for injections (2 mg/mL) was prepared in 200 mM ß-Cyclodextrin sulfobutyl ether/PBS. To completely dissolve the compound, the vial was first placed into a sonication bath (30 min, 37°C) and then heated up in a 98 °C water bath for 30 min. Similarly, the solution of DKFZ-682 (4 mg/mL) was prepared. Cisplatin and etoposide were dissolved in 60 mM ß-Cyclodextrin sulfobutyl ether/PBS at a concentration of 0.3 mg/mL and 0.75 mg/mL, respectively. CDDO-Me was first dissolved in DMSO to make a stock at a concentration of 20 mg/mL. This stock was then mixed with PEG300 (40 %), Tween-80 (5 %) and PBS (50 %) to dilute it to 1 mg/mL. Female mice (6–7 weeks old) of the nude strain BALB/c (Charles River) were used for the dose escalation study and the repeated-dose efficacy study with xenografts. To determine the maximum tolerated dose (MTD) of DKFZ-608 for intraperitoneal administration, three animals were injected daily, starting from 5 mg/kg, with doses of DKFZ-608 increasing every three days to 7.5, 10, 12.5, 15 and finally 20 mg/kg. Body weight was monitored daily and weight loss was observed only at a dose of 20 mg/kg. Therefore, we defined the MTD as 15 mg/kg, which is the dose that was used for tumor treatment. To generate a subcutaneous SCLC-tumor model, 3 x 10^7^ H209 cells in 100 µL of ice-cold PBS were injected subcutaneously into the flank. 6 weeks later tumor bearing animals (tumor size 4–8 mm) were selected and randomly distributed to experimental groups of 10 animals each. Tested substances were administered for 3 weeks (chemotherapy) or 40 days (DKFZ-608). Specifically, 0.2 ml of each drug solution was injected per 20 g mouse to obtain a dose of 5–20 mg/kg DKFZ-608 (daily *i.p.* injections), 3 mg/kg cisplatin (on Monday) and 7.5 mg/kg etoposide (on Wednesday and Friday). Tumor size was determined twice a week in two dimension using calipers and tumor volumes were calculated by the formula (width^2^ × length)/2.

Female mice (8–10 weeks old) of the immunodeficient strain NSG were used for the dose escalation study of DKFZ-682 in combination with CDDO-Me and the dose efficacy study with xenografts. To determine the MTD of DKFZ-682 for intraperitoneal administration, three animals were injected daily, starting from 4 mg/kg, with doses of DKFZ-682 increasing every 1–3 days to 6, 8, 10 and finally 12 mg/kg. At the highest dose, the animals reached the human endpoint. Body weight was monitored daily and weight loss did not exceed 20 % at a dose of 10 mg/kg. Therefore, we defined the MTD of DKFZ-682 upon CDDO-Me co-treatment as 10 mg/kg. To generate a subcutaneous SCLC- tumor model, H526 cells were resuspended in a 1:1 (vol/vol) mix of ice-cold growth factor-reduced Matrigel (Corning) and PBS. Overall, 100 µl of this cell suspension containing 2×10^6^ H526 cells was injected subcutaneously into the flank of anesthetized mice. After detection of palpable tumors 2 weeks later, tumor bearing animals were randomly distributed to experimental groups. Tested substances were administered via intraperitoneal injection daily for 3 weeks. Tumor size was determined every 2–3 days using calipers and tumor volumes were calculated by the formula (width^2^ × length)/2. CDDO-Me was used at a concentration of 3 mg/kg, DKFZ-682 at 4 mg/kg and 10 mg/kg alone and in combination with CDDO-Me, respectively.

To evaluate organ damage markers, the animals were sacrificed by isoflurane overdose and blood was collected from the portal vein using 24G needles (BD Microlance 3) into microtubes with clot activator (Microvette 500 CAT-Gel, Saarstedt). After incubation for 30 min at room temperature, blood samples were centrifuged for 5 min at 10,000 *g* at room temperature and the upper cell-free layer consisting of serum was transferred into a new tube and stored at −80 °C until further analysis. Pieces of tissues (liver, kidney, lung, heart, tumor) were collected for further protein (dry) and transcript (into RNAlater, Sigma-Aldrich) analysis and stored at −80 °C until further processing.

### Organ damage markers

Kidney damage was quantified as the level of urea in serum using the Urea Nitrogen (BUN) Colorimetric Detection Kit (Invitrogen) according to the manufacturer’s instructions, adding 4 uL of serum per well. As proxies for liver damage, the activities of alanine aminotransferase (ALT) and aspartate aminotransferase (AST) were measured in serum samples using Alanine Aminotransferase Activity Assay Kit (MAK052, Sigma-Aldrich) and Aspartate Aminotransferase Activity Assay Kit (MAK055, Sigma-Aldrich), respectively, according to the manufacturer’s instructions and adding 3 uL and 5 uL of serum per well for ALT and AST activity, respectively.

### Flow cytometry

As an indicator of ferroptosis, lipid peroxidation was analyzed in cells stained with Bodipy 581/591 C11 (Invitrogen). Cells were stained with 3 µM Bodipy 581/591 C11 diluted in the culture medium for 30 min at 37 °C. Afterwards, cells were washed with PBS, and analyzed using the flow cytometer BD LSR Fortessa (BD Biosciences). The ratio of the oxidized (excitation 488 nm, emission 530/30) and reduced (excitation 561 nm, emission 610/20) dye was calculated for each cell in the FlowJo software.

For detection of ROS/RNS level 150,000 cells/well/1 mL (adherent cells) or 500,000 cells/mL (suspension cells) were seeded in 12-well plate. The next day, cells were treated with DMSO (unstained control), 5 µM CM-H_2_DCFDA (Invitrogen, Thermo Scientific, Cat No C6827), 1:250 OxiVision Green peroxide sensor (AAT Bioquest, Cat No. 11506, powder was solved in 200 µL DMSO) or 5 µM DAF-FM Diacetate (Invitrogen, Thermo Scientific, Cat No. D23844) at 37 °C for 30 min; 1:400 DAX-J2™ PON Green (AAT Bioquest, Cat No. 16317) at 37 °C for 60 min. Then cells were detached with trypsin (adherent cells) or washed (suspension cells), spun down 5 min at 1200 rpm and finally each cell pellet was resuspended in 500 µL PBS with 1 % FBS and analyzed by the flow cytometer Guava easyCyte 14HT (Luminex). The fluorescence of all above-mentioned dyes was analyzed in the Green-B channel (excitation 488 nm, emission 512/18 nm) while counting 10,000 cells per sample. For ROS Brite 570 (10 µM, AAT Bioquest, Cat No. 16000) staining, cells were first detached with trypsin (adherent cells) or washed (suspension cells) and then incubated with ROS Brite 570 at 37 °C. After 20 min cells were spun down 5 min at 130 *g* and cell pellet was resuspended in 500 µL PBS with 1 % FCS and analyzed by flow cytometry (Yellow-G channel: excitation 532 nm, emission 575/25).

Treatment: cells were co-incubated with OxiVision Green peroxide sensor and drug; pre-incubated with drug, then detached with trypsin (adherent cells) or spun down (suspension cells), washed and further incubated with ROS Brite 570. Stained but untreated cells were used as control.

### Measurement of ROS levels in cytoplasm and mitochondria

The effect of drugs on roGFP2-Orp1 oxidation was quantified as described previously [15].

### Metabolomic analysis of nucleotide intermediates

To analyze nucleotide intermediates (NADP^+^ and NADPH), adherent cells (HaCaT, Beas-2B) were quickly rinsed with ice-cold saline (0.9 % NaCl) and scraped off into 1 mL of ice-cold saline. Cell suspension was transferred into a microtube and spun down at 500 g for 1 min at 4 °C. After removal of supernatant, cell pellets were snap-frozen in liquid nitrogen and stored at −80 °C until further processing. Suspension cells were first pelleted at 500 g for 2 min at 4 °C in 15-mL tubes. Pellets were resuspended in 1 mL of ice-cold saline, cell suspension was transferred into a microtube and spun down at 500 g for 2 min at 4 °C. After removal of supernatant, cell pellets were snap-frozen in liquid nitrogen and stored at −80 °C until further processing. Metabolites were extracted and LC-MS/MS analysis performed by the Metabolomics Core Technology Platform (Heidelberg). After extraction, pelleted cell debris was lysed in Pierce^TM^ RIPA buffer (Thermo Scientific, 89900) supplemented with a protease inhibitor cocktail (Serva) and sonicated on ice for 10 s using a probe sonicator (10 % amplitude). The protein concentration in lysates was measured using Pierce BCA Protein Assay Kit (Thermo Scientific) with BSA as a standard. Metabolite results were expressed as unitless peak area corrected with d5-tryp as an internal standard for batch effect correction. The data were normalized to protein amount.

### Detection of glutathione

Intracellular GSH levels were quantified using the Reduced Glutathione Assay Kit (Abcam, ab235670) following the manufacturer’s instructions. To prepare the samples, adherent cells (HaCaT, Beas-2B) were detached using trypsin, washed first with culture medium to neutralize trypsin activity, and then washed with PBS. One million pelleted cells were resuspended in 30 µL of 5 % sulfosalicylic acid (SSA) solution and incubated on ice for 10 min. After centrifugation at 12,000 g at 4°C for 10 min, the supernatant was collected as the sample solution. The samples were then diluted 20-fold with GSH Assay Buffer, and 10 µL of the diluted samples were added to the wells of a black 96-well plate. The sample volumes were adjusted to 20 µL with GSH Assay Buffer, followed by the addition of 80 µL of Reaction Mix to each well. Fluorescence was measured in kinetic mode at room temperature for 60 min with 2-minute intervals in a CLARIOstar plate reader (BMG LABTECH).

Intracellular levels of glutathione (GSH and GSSG) in the cells were determined using the GSH/GSSG-Glo^TM^ Assay (Promega, V6611). Cells (HaCaT and Beas-2B: 2 × 10⁵ cells in 2 mL of medium per well in a 6-well plate; H82 and H69: 10⁶ cells in 2 mL of medium per well in a 12-well plate) were seeded. The following day, the test compound was added to each well and incubated for 24 h. Then, the cells were harvested with lysis buffer (0.5% NP-40 in PBS) with or without N-ethylmaleimide (NEM, 2.5 mM), and glutathione levels were quantified using the GSH/GSSG-Glo™ Assay according to the manufacturer’s instructions. Luminescence was measured using a FLUOstar OPTIMA plate reader (BMG LABTECH).

### Hoechst 33342/propidium iodide double staining

The cells seeded in a 96-well plate were incubated with Hoechst 33342 (1 µg/mL) and propidium iodide (1 µg/mL) in a dark incubator at 37 °C for 30 min. Images were obtained using a Leica fluorescence microscope (40x objective magnification) combined with a digital camera and Las Ez (version 3.1.0) software.

### Software and statistical analysis

All figures and statistical analyses were generated using Affinity Designer and GraphPad Prism 9 software. Schematic overview figures were created with BioRender.com.

Data are presented as mean ± SD. For the comparison of protein levels and EC_50_ values, a two-tailed unpaired *t*-test was used. For the comparison of expression values, an unpaired multiple t-test, comparing DMSO versus drug treatment, was used.

## Data availability

The RNA sequencing and microarray data generated in this study was deposited in the NCBI Gene Expression Omnibus (GEO) database under the primary accession number GSE279100. (Token for access will be made available upon request).

The publicly available data used in this study are available in the European Genome-Phenome Archive under the primary accession number EGAS00001000334 and in [75, 76]

The remaining data are available within the Article, Supplementary Information or Source Data file.

---

**Supplementary Table S1:** Chemical compounds.

| Chemical compound | Company | Catalog No |
| --- | --- | --- |
| 1S,3R-RSL 3 (RSL3) | Sigma-Aldrich | SML2234 |
| Auranofin | Sigma-Aldrich | A6733 |
| Azacitidine (AZA) | Selleckchem | S1782 |
| Beta-Cyclodextrin sulfobutyl ether sodium salt | Biosynth | OC15979 |
| 2-Hydroxy β-Cyclodextrin | Sigma-Aldrich | 332607 |
| Buthionine sulfoximine (BSO) | Sigma-Aldrich | B2640 |
| CDDO-Me | Tocris | 6646 |
| Cisplatin | Santa Cruz Biotechnology | SC-200896A |
| CPUY192018 (CPUY) | Biotrend | AOB9974-1 |
| Decitabine (DAC) | Hözel Diagnostica | TMO-T1508 |
| Diamide (DA) | Sigma-Aldrich | D3648 |
| Dimethylsulfoxide (DMSO) | Sigma-Aldrich | D2650 |
| Deferoxamine (DFO) | Sigma-Aldrich | D9533 |
| DKFZ-608 | produced by AG Gunkel/Miller, DKFZ |  |
| DKFZ-682 | produced by AG Gunkel/Miller, DKFZ |  |
| Dimethyl fumarate (DMF) | Sigma-Aldrich | 242926 |
| Dithiothreitol (DTT) | Serva | 20711.02 |
| Ferostatin-1 (Fer-1) | Sigma-Aldrich | SML0583 |
| G6PDi-1 | Sigma-Aldrich | SML2980 |
| Hoechst 33342 | Cayman chemical | 15547 |
| Necrostatin (Nec-1) | Merck | 480065 |
| Olaparib (AZD2281) | Cell Signaling | 93852 |
| Propidium Iodide (PI) | Sigma-Aldrich | P4864 |
| Puromycin | Sigma-Aldrich | P9620 |
| Sodium aurothiomalate | Fisher Scientific | 11344707 |
| Tetraethyldithiuram disulphide | Sigma-Aldrich | 86720 |
| Z-VAD-FMK | Adooq Bioscience | A12373 |
| TRFS green | produced by AG Gunkel/Miller, DKFZ, as described by Zhang et al. [71] |  |

**Supplementary Table S2:**
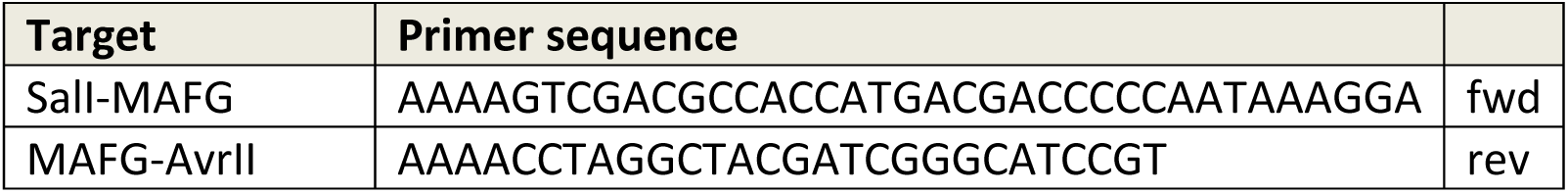
Primers Cloning ORFs into pLIX-403.

**Supplementary Table S3:** Primary and secondary antibodies.

| Antigen | Product / Company | Species | Dilution WB |
| --- | --- | --- | --- |
| AIF | 4642/ Cell Signaling | rabbit | 1:1000 |
| AIFM2/FSP1 | 20886-1-AP/Proteintech | rabbit | 1:1000 |
| AKR1C3 | A6229/ Sigma-Aldrich | mouse | 1:500 |
| BACH1 | NBP1-71814/ Novus | rabbit | 1:2000 |
| FTL | ab69090/ Abcam | rabbit | 1:1000 |
| GAPDH | sc-365062/ Santa Cruz | mouse | 1:1000 |
| GCLC | ab207777/ Abcam | rabbit | 1:1000 |
| GCLM | HPA023696/ Sigma-Aldrich | rabbit | 1:100 |
| GPX2 | ab137431/ Abcam | rabbit | 1:1000 |
| GSR | ab124995/ Abcam | rabbit | 1:1000 |
| Lamin A/C | 2032/ Cell Signaling | rabbit | 1:1000 |
| MAFG | ab154318/ Abcam | rabbit | 1:1000 |
| ME1 | ab97445/ Abcam | rabbit | 1:1000 |
| NQO1 | ab28947/ Abcam | mouse | 1:1000 |
| NRF2 | ab62352/ Abcam | rabbit | 1:1000 |
| PRDX1 | 8499/ Cell Signaling | rabbit | 1:1000 |
| PRDX3 | ab128953/ Abcam | rabbit | 1:2000 |
| SLC7A11 | 12691/ Cell Signaling | rabbit | 1:1000 |
| TXN (TRX1) | ab133524/ Abcam | rabbit | 1:10000 |
| TXNRD1 (TRXR1) | ab16820/ Abcam | rabbit | 1:1000 |
| β-Tubulin | T0198/ Sigma-Aldrich | mouse | 1:1000 |
| IRDye 680LT anti-mouse IgG | 926-68022/ LI-COR | donkey | 1:5000 |
| RDye 680LT anti-rabbit IgG | 926-68023/ LI-COR | donkey | 1:5000 |
| StarBright Blue 520 anti-rabbit IgG | 12005869/ Biorad | goat | 1:3000 |
| StarBright Blue 700 anti-rabbit IgG | 12004162/ Biorad | goat | 1:3000 |
| StarBright Blue 520 anti-mouse IgG | 12005866/ Biorad | goat | 1:3000 |
| StarBright Blue 700 anti-mouse IgG | 12004159/ Biorad | goat | 1:3000 |

**Supplementary Table S4:**
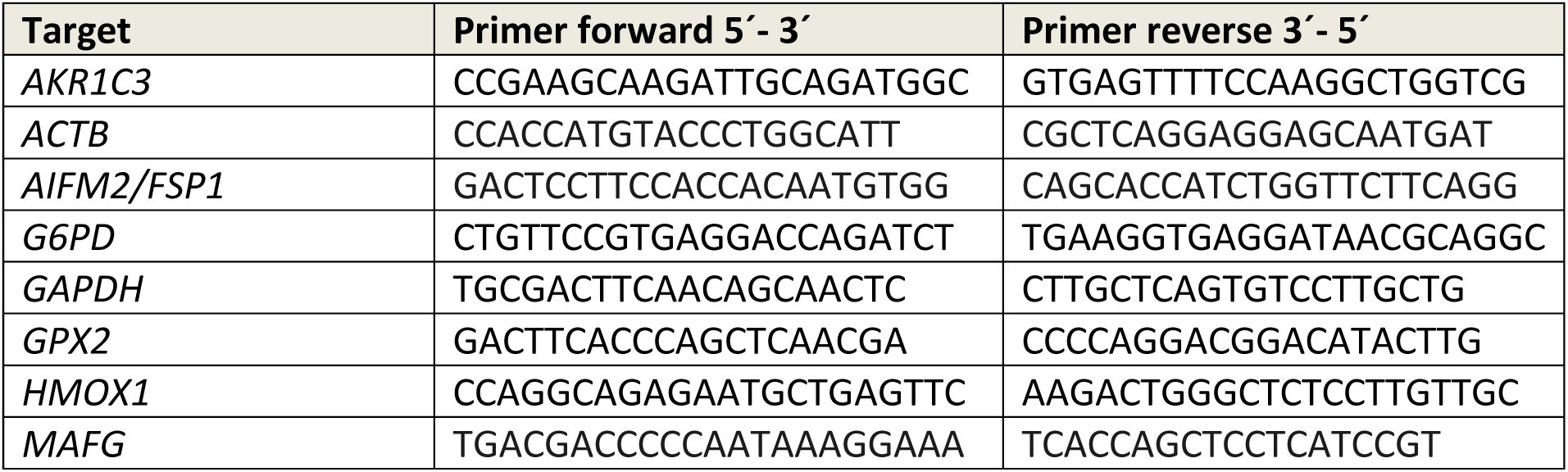
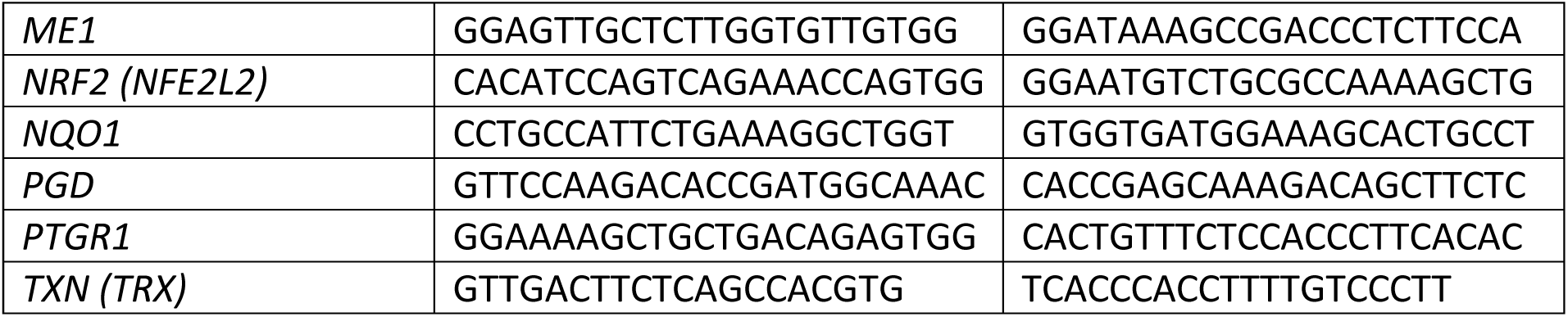
qPCR primers Human.

## Acknowledgments

We thank the Helmholtz Validation Fund (grant number HVF-0069) and Else Kröner-Fresenius-Stiftung Translationsförderung (EKFS-Theropo) for supporting this work. M. Muckenthaler acknowledges support from the Deutsche Forschungsgemeinschaft (FerrOs-FOR5146; GRK2727; SPP2306). Hamed Alborzinia acknowledges the support of DFG priority program SPP230. Moritz Mall acknowledges support by funding from CellNetworks (EXC81), ERC StG No 804710, and the Hector Stiftung II GmbH. Fellowships were provided by the Helmholtz International Graduate School to B.L.. We thank the Microarray Core Facility of the German Cancer Research Center (DKFZ) for providing excellent Expression Profiling services and the Illumina methylation arrays and related services. We thank the NGS Core Facility of the German Cancer Research Center (DKFZ) for providing excellent RNAseq services. We also thank Jonas Gross, Tobias Herrmann, Nawid Albinger and Jonas Kolibius for the valuable work during their lab internship, which helped to build a robust understanding of the mode of action of DKFZ-608.

## Author contributions

Jana Samarin, Hana Nuskova, Piotr Fabrowski, Mona Malz, Eberhard Amtmann, Minerva J. Taeubert, Daniel Pastor-Flores and Daniel Kazdal have made substantial contributions to the acquisition, analysis, or interpretation of data. Roman Kurilov and Julia Frede have substantially contributed by bioinformatics analysis of data and interpretations. Nicole de Vries, Hannelore Pink, Franziska Deis, Kamini Kaushal, Bryce Lim, Taishi Kanamaru and Glynis Klinke made substantial contributions to the acquisition of data. Gerhard Hamilton was instrumental for data obtained with circulating tumor cells. Moritz Mall and Tobias P. Dick made substantial contributions to the interpretation of data. Martina Muckenthaler and Martin Sos have substantially contributed to drafting the work and substantively revised it. Aubry K. Miller, Hamed Alborzinia and Nikolas Gunkel have made substantial contributions to the conception and design of the work and have drafted and substantively revised it. Aubry K. Miller was instrumental for the design and synthesis of TXNRD1 inhibitors.

